# The darkest place is under the candlestick - healthy urogenital tract as a source of UTI-related *Escherichia coli* lineages

**DOI:** 10.1101/2021.08.06.455400

**Authors:** Magdalena Ksiezarek, Angela Novais, Luisa Peixe

## Abstract

Since the discovery of the urinary microbiome, including identification of *Escherichia coli* in healthy hosts, its involvement in UTI development is a subject of high interest.

We explored population diversity and antimicrobial resistance of *E. coli* from urogenital microbiome of asymptomatic and recurrent UTI (rUTI) women. We also evaluated the genomic relationship between extraintestinal pathogenic *E. coli* (ExPEC) strains from healthy and diseased hosts, particularly of the ST131 lineage.

*E. coli* was highly prevalent in asymptomatic women (48%) with slightly higher prevalence in vaginal samples comparing to urine, and occasionally with multiclonal population in the same individual. B2 was the most frequent phylogenetic group, with most strains classified as ExPEC. We demonstrated that virulence associated genes profile does not allow to distinguish strains isolated from healthy and rUTI host. We identified *E. coli* widespread lineages e.g., sequence types (ST) 127, ST131 (asymptomatic cohort) and ST73, ST131 (rUTI), frequently resistant to at least one antibiotic. Phylogenomics of ST131 and other ExPEC lineages revealed close relatedness between healthy and diseased host.

These findings demonstrate that healthy urogenital microbiome is a source of potentially pathogenic and antibiotic resistant *E. coli* strains, including globally spread *E. coli* lineages causing UTI including ST131.

## INTRODUCTION

Urinary tract infections (UTIs) are reported as the most common infections worldwide, that occur more frequently in women than men due to anatomical differences, and its incidence increases with age or sexual activity^1–4^. Although several bacterial species are reported as causative agents of UTIs (e.g., *Escherichia coli*, *Klebsiella* sp., *Proteus* sp., *Staphylococcus saprophyticus*), *E. coli* is responsible for the majority (up to 75%) of both uncomplicated and complicated UTIs^2, 5–7^.

Extraintestinal pathogenic *E. coli* (ExPEC) strains are mostly responsible for these infections, which possess genetic characteristics that seem to favor pathogenicity of particular strains but also *E. coli* persistence in the urinary tract^8–10^. Indeed, most UTIs are caused by a subset of ExPEC strains [e.g., sequence type (ST) 131, ST95, ST69, ST73, ST127, ST12] that are highly disseminated across different continents and populations, that represent great clinical challenges not only because they are often resistant to main therapeutic choices but also because of their extended reservoirs (human and animal intestines and the environment)^11–16^.

To date, multiple studies evaluated putative human-related reservoirs of ExPEC strains causing UTI. The human gut or vagina colonization with certain strains of *E. coli* has been considered a risk factor for developing UTI^17–21^.

Additionally, the existence of intracellular bacterial communities or quiescent intracellular reservoirs that may silently persist within epithelial cells of urogenital tract also contribute to recurrent UTI (rUTI) development^7, 22^.

On the other hand, the recently disproven urine sterility paradigm has challenged our understanding on urogenital tract health, and introduced the possibility of a urinary/urogenital reservoir for UTI pathogens^23, 24^. With the identification of the female urogenital microbiome (FUM)^25^ and later the urinary microbiome^26^, the origin of *E. coli* causing UTIs was re-evaluated. Studies comparing FUM composition of female healthy and UTI cohort suggested that the urinary tract may harbor putative uropathogenic bacterial species, including *E. coli*^27, 28^. Although it is not among the most prevalent bacterial species, *E. coli* is also part of FUM identified in voided or catheterized urine of healthy women^26, 29–31^. Moreover, a high inter-individual variability in *E. coli* relative abundance (varying from 10 to >10^5 CFU/ml) in healthy population was reported^27, 29, 31^. Thus, since *E. coli* is the most frequent pathogen causing UTI and it can be found in healthy FUM (sometimes in high relative abundance), the possibility of the urinary tract itself acting as a source of ExPEC lineages causing UTIs is of utmost relevance.

Till now, the urinary research focused mostly on taxonomic profiling of FUM structure, reaching at its best the species level, and only a few studies investigating healthy microbiome population at the strain level are available. Thomas-White and co-workers^28^ have shown urinary and vaginal interconnection for potentially pathogenic and health promoting species, based on strains similarity. Although of great relevance, this study did not explore any further gene content and specific virulence potential of *E. coli* strains. The genomic strain level characterization was provided in a recent study from Garretto *et al*., which evaluated the association of genomic content of urinary isolates and the presence of lower urinary tract symptoms^32^. Importantly, Garretto *et al*., concluded that there is no specificity regarding *E. coli* abundance, gene content, neither the composition of microbiome itself that could predict UTI status. Of note, this study focused exclusively on urinary isolates and only 6 strains from investigated collection originated from asymptomatic women^32^. Thus, to date, strains colonizing urogenital tract of healthy women have been scarcely characterized.

In our study, we aimed to analyze prevalence, population diversity and antimicrobial resistance of *E. coli* isolated from urine and vaginal samples of healthy asymptomatic women and women with history of rUTI. We characterized *E. coli* at the strain level and performed thorough analysis of their phylogenetic background and virulome. Furthermore, we provided an in-depth phylogenomic analysis of B2 ExPEC strains, antimicrobial resistance and plasmid profiling. We also performed comparative genomics for ST131 *E. coli* colonizing the urogenital tract of women with different health conditions with *E. coli* genomes from human origin available in public databases.

## RESULTS

### Frequency and diversity of *E. coli* in urinary and vaginal microbiome

*E. coli* isolates were identified in 23 samples from a total of 64 analyzed samples (31 voided urine and 30 vaginal swabs from healthy cohort, 3 voided urine samples from rUTI-cohort). They were identified in 48% (10/21) of healthy donors (29% of urine samples and 37% of vaginal samples) in loads ranging from 1×10 CFU/ml – 1×10^7 CFU/ml (median 1×10 CFU/ml), and in all rUTI urine samples in loads ranging from 1×10 - 1×10^4 CFU/ml.

A set of 25 *E. coli* isolates representing unique virulence factor (VF) profiles from diverse phylogenetic groups per different sample type were preliminarily selected. They were recovered from 9 urine and 11 vaginal samples from healthy donors and 3 urine samples from rUTI cohort, for which information on donor, sample type, phylogenetic group, clones’ similarity by pulsed-field gel electrophoresis (PFGE) and ExPEC status is compiled in **Supplementary Table S1**. Interestingly, 2 donors carried different *E. coli* strains in urine and vaginal samples and 2 donors had diverse *E. coli* in urine and/or vagina.

*E. coli* phylogenetic group B2 isolates (56%) were more frequently detected, followed by F (16%) and D (12%). Remaining phylogenetic groups (A, C and E) were detected occasionally (4-8%) (Table 1). It is of interest to highlight that a statistically significant (p<0.05) association was found for phylogenetic group B2 and isolates obtained from rUTI women, whereas *E. coli* isolates from healthy women belonged to different phylogenetic groups (11 B2, 2A, 1 C, 3D, 1E, 4F; Table 1). Some VFs were found significantly associated with rUTI *E. coli* strains (p<0.05), such as adhesin *yfcV*, protectin *traT*, miscellaneous *usp*, *pafP* and *upaH* (Table 1). There was no significant association between phylogenetic groups distribution or VF profiles and the origin (urine or vaginal) of the sample.

**Table 1.** Detection and prevalence (%) of phylogenetic groups and 41 putative virulence genes among 25 unique *E. coli* strains per each sample type/cohort. Only p values of <0.05 are presented in the table in bold font. There was no isolate positive for papG allele II, bmaE, gafD. None of the isolates tested had two copies of hlyA operon. None of phylogenetic group or virulence factors was detected as significantly different between urinary and vaginal isolates.

|  | Total<br>(n=25) | H urine<br>(n=10) | H vaginal<br>(n=12) | rUTI urine<br>(n=3) | p value of H<br>urine vs rUTI<br>urine | B2 (n=14) | non-B2<br>(n=11) | p value of B2<br>vs non-B2 |
| --- | --- | --- | --- | --- | --- | --- | --- | --- |
| **Phylogenetic**<br>**group** |  |  |  |  |  |  |  |  |
| A | 2 (8) | 1 (10) | 1 (8) | 0 |  |  |  |  |
| B2 | 14 (56) | 4 (40) | 7 (58) | 3 (100) | **<0.01** |  |  |  |
| C | 1 (4) | 0 | 1 (8) | 0 |  |  |  |  |
| D | 3 (12) | 2 (20) | 1 (8) | 0 |  |  |  |  |
| E | 1 (4) | 1 (10) | 0 | 0 |  |  |  |  |
| F | 4 (16) | 2 (20) | 2 (17) | 0 |  |  |  |  |
| **Adhesins** |  |  |  |  |  |  |  |  |
| *fimH* | 23 (92) | 9 (90) | 11 (92) | 3 (100) |  | 14 (100) | 9 (82) |  |
| *papAH* | 4 (16) | 2 (20) | 1 (8) | 1 (33) |  | 2 (14) | 2 (18) |  |
| *papC* | 8 (32) | 3 (30) | 4 (33) | 1 (33) |  | 4 (29) | 4 (36) |  |
| *papEF* | 8 (32) | 3 (30) | 4 (33) | 1 (33) |  | 3 (21) | 5 (45) |  |
| *papG II,III* | 2 (8) | 0 | 1 (8) | 1 (33) |  | 2 (14) | 0 |  |
| *papG allele III* | 4 (16) | 1 (10) | 2 (17) | 1 (33) |  | 4 (29) | 0 | **<0.05** |
| *sfa/focD* | 8 (32) | 2 (20) | 4 (33) | 2 (67) |  | 8 (57) | 0 | **<0.01** |
| *sfaS* | 2 (8) | 0 | 1 (8) | 1 (33) |  | 2 (14) | 0 |  |
| *focG* | 7 (28) | 2 (20) | 4 (33) | 1 (33) |  | 7 (50) | 0 | **<0.01** |
| *afa/draBC* | 3 (12) | 2 (20) | 1 (8) | 0 |  | 2 (14) | 1 (9) |  |
| *iha* | 8 (32) | 4 (40) | 3 (25) | 1 (33) |  | 4 (29) | 4 (36) |  |
| *matB* | 21 (84) | 8 (80) | 10 (83) | 3 (100) |  | 14 (100) | 7 (64) | **<0.05** |
| *yfcV* | 18 (72) | 6 (60) | 9 (75) | 3 (100) | **<0.05** | 14 (100) | 4 (36) | **<0.01** |
| **Toxins** |  |  |  |  |  |  |  |  |
| *hlyA* | 5 (20) | 1 (10) | 2 (17) | 2 (67) |  | 5 (36) | 0 | **<0.05** |
| *cnfI* | 5 (20) | 1 (10) | 2 (17) | 2 (67) |  | 5 (36) | 0 | **<0.05** |
| *sat* | 10 (40) | 4 (40) | 5 (42) | 1 (33) |  | 4 (29) | 6 (55) |  |
| *tsh* | 10 (40) | 3 (30) | 5 (42) | 2 (67) |  | 8 (57) | 2 (18) | **<0.05** |
| *vat* | 7 (28) | 1 (10) | 4 (33) | 2 (67) |  | 7 (50) | 0 | **<0.01** |
| *tosA* | 5 (20) | 2 (20) | 2 (17) | 1 (33) |  | 5 (36) | 0 | **<0.05** |
| **Siderophores** |  |  |  |  |  |  |  |  |
| *fyuA* | 17 (68) | 8 (80) | 6 (50) | 3 (100) |  | 12 (86) | 5 (45) | **<0.05** |
| *iutA* | 9 (36) | 5 (50) | 3 (25) | 1 (33) |  | 5 (36) | 4 (36) |  |
| *iroN* | 10 (40) | 3 (30) | 5 (42) | 2 (67) |  | 9 (64) | 1 (9) | **<0.01** |
| *ireA* | 3 (12) | 1 (10) | 2 (17) | 0 |  | 2 (14) | 1 (9) |  |
| **Capsule** |  |  |  |  |  |  |  |  |
| *kpsMT II* | 14 (56) | 5 (50) | 7 (58) | 2 (67) |  | 11 (79) | 3 (27) | **<0.05** |
| *kpsMT III* | 3 (12) | 1 (10) | 1 (8) | 1 (33) |  | 1 (7) | 2 (18) |  |
| *kpsMT K1* | 5 (20) | 1 (10) | 4 (33) | 0 |  | 3 (21) | 2 (18) |  |
| *kpsMT K5* | 7 (28) | 3 (30) | 3 (25) | 1 (33) |  | 7 (50) | 0 | **<0.01** |
| **Protectins** |  |  |  |  |  |  |  |  |
| *rfc* | 1 (4) | 0 | 0 | 1 (33) |  | 1 (7) | 0 |  |
| *cvaC* | 2 (8) | 1 (10) | 1 (8) | 0 |  | 1 (7) | 1 (9) |  |
| *traT* | 15 (60) | 6 (60) | 6 (50) | 3 (100) | **<0.05** | 7 (50) | 8 (73) |  |
| *iss* | 2 (8) | 1 (10) | 1 (8) | 0 |  | 1 (7) | 1 (9) |  |
| **Invasins** |  |  |  |  |  |  |  |  |
| *ibeA* | 3 (12) | 0 | 3 (25) | 0 |  | 3 (21) | 0 |  |
| *gimB* | 2 (8) | 0 | 1 (8) | 1 (33) |  | 2 (14) | 0 |  |
| **Miscellaneous** |  |  |  |  |  |  |  |  |
| *usp* | 16 (64) | 5 (50) | 8 (67) | 3 (100) | **<0.05** | 12 (86) | 4 (36) | **<0.05** |
| *ompT* | 21 (84) | 8 (80) | 10 (83) | 3 (100) |  | 14 (100) | 7 (64) | **<0.05** |
| *PAI (malX)* | 15 (60) | 7 (70) | 5 (42) | 3 (100) |  | 12 (86) | 3 (27) | **<0.01** |
| *pafP* | 18 (72) | 6 (60) | 9 (75) | 3 (100) | **<0.05** | 14 (100) | 4 (36) | **<0.01** |
| *upaH* | 14 (56) | 4 (40) | 7 (58) | 3 (100) | **<0.01** | 14 (100) | 0 | **<0.01** |

Certain VFs were overall highly prevalent such as adhesins *fimH* (92%), *matB* (84%) or *ompT* (84%). The siderophore *fyuA* (68%) or protectin *traT* (60%) were also frequent. We also found that many strains possess *kpsMT II* type (56%).

As expected according to results from other *E. coli* series, several adhesins (e.g., *focG*, *matB*, *yfcV*), most toxins (e.g., *hlyA*, *cnf1*, *vat*), siderophores (*fyuA*, *iroN*), *kpsMT II* and K5 capsule type and miscellaneous (e.g., *usp*, *ompT*, *malX*) were significantly enriched in strains of phylogenetic group B2 (Table 1). All but one B2 strains were classified as ExPEC. Furthermore, two isolates belonging to phylogenetic group F isolated from urine of healthy donors (c1Ub_48 and c10Ua_105) were also considered ExPEC (**Supplementary Table S1**).

### Virulence profile characterization of urogenital *E. coli*

A detailed analysis of VF profiles observed for representative strains and phylogenetic groups per individual was performed by hierarchical clustering analysis (Figure 1). Vaginal isolates that presented identical PFGE profiles to urine isolates from the same individual and phylogenetic group were discarded from this analysis (**Supplementary Figure S1**). Based on this strategy, we selected 19 unique *E. coli* strains identified from healthy women and women with rUTI, independently on their sample type (**Supplementary Table S1**).

**Figure 1.**
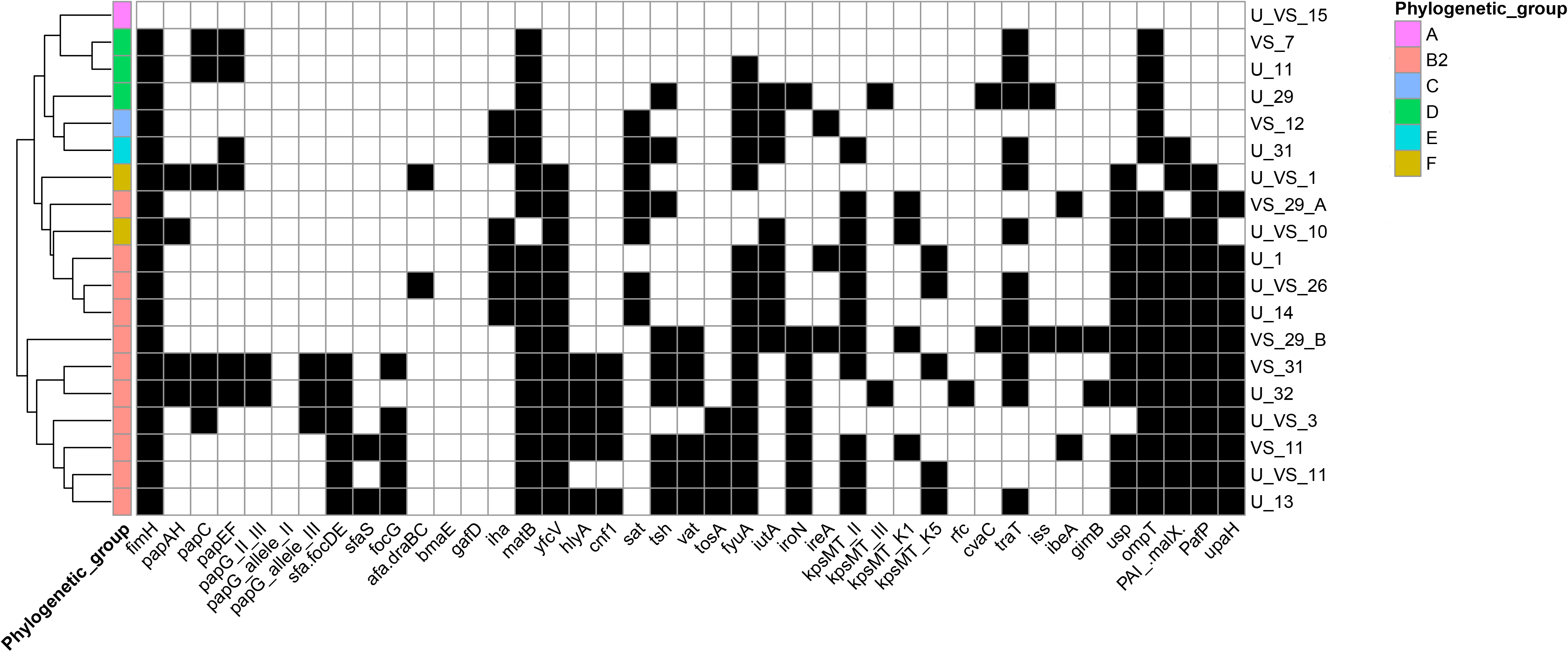
Heatmap representing presence/absence matrix on putative virulence genes among urogenital *E. coli* collection. Left dendrogram represents clustering of isolates based on Euclidean distance. Detected virulence genes are represented with black squares within the heatmap, while white stands for absence. Right list of unique *E. coli* profiles is coded with U and/or VS at the begin which stands for urine and/or vaginal swab, respectively. The following number within the code stands for donor. Phylogenetic groups are encoded in the left colorful bar, according to the legend.

A hierarchical clustering analysis of VF profiles of representative *E. coli* strains showed that isolates belonging to B2 phylogenetic group were divided into two clusters (Figure 1). One of which included most of B2 isolates (7/11; 64%) and was enriched in adhesins belonging to *pap* operon and various toxins i.e., *hlyA*, *cnf1*, *tsh*, *vat* and *tosA*. The remaining B2 isolates (4/11) clustered with phylogenetic group F ExPEC strains. The two clusters observed contain isolates from mixed cohorts i.e., clustering based on VF profile does not allow to distinguish strains from healthy or rUTI host.

### Genomic characterization and antimicrobial resistance of B2 strains

Eleven B2 ExPEC strains were further selected for whole genome sequencing, phylogenetic analysis and comparative genomics. Strains characterization, characteristics of draft genomes assemblies and accession numbers are shown in Table 2. B2 isolates from healthy donors were identified as ST127, ST131, ST140 (clonal complex 95), ST452, ST569, ST681, ST998 and ST1154 (clonal complex 73), while isolates from women with history of rUTI belonged to ST12, ST73 and ST131, some of which are recognized widespread *E. coli* lineages (e.g., ST131, ST127, ST73). The strains represented various serogroups, including O2, O6, O8, O25, O46 and O81 among isolates from healthy hosts, and O4, O15 and O22 among isolates from rUTI cohort (Table 2).

**Table 2.** Details on strains origin, characteristics of draft genome assemblies (statistics for >500 bp contigs) and accession numbers for 11 urogenital *E. coli* strains. ST - sequence type; SLV - single locus variant.

| strain | origin | host | ST | SL<br>V | serotype | total<br>length | nr<br>contigs | N50 | Accession | BioSample |
| --- | --- | --- | --- | --- | --- | --- | --- | --- | --- | --- |
| c1Ub_56_AE | urine | healthy | 452 | - | O81:H27 | 5294227 | 188 | 128712 | JAECYT000000000 | SAMN17025145 |
| c3Ub_1_AE | urine | healthy | 681 | - | O8:H10 | 5286427 | 59 | 348018 | JAECYS000000000 | SAMN17025146 |
| c11Ub_17_AN | urine | healthy | 1154 | 73 | O2:H1 | 4908002 | 32 | 625852 | JAECYR000000000 | SAMN17025147 |
| c11VSb_12_AN | vagina | healthy | 998 | - | O2:H6 | 5075113 | 29 | 510981 | JAECYQ000000000 | SAMN17025148 |
| c13Ua_2_AN | urine | rUTI | 73 | - | O22:H1 | 5170382 | 84 | 320578 | JAECYP000000000 | SAMN17025149 |
| c14Ub_22_AE | urine | rUTI | 131 | - | O15:H5 | 5049786 | 76 | 323647 | JAECYO000000000 | SAMN17025150 |
| c26Ub_7_AN | urine | healthy | 131 | - | O25:H4 | 5170043 | 61 | 305535 | JAECYN000000000 | SAMN17025151 |
| c29VSa_23_AE | vagina | healthy | 569 | - | O46:H31 | 4863018 | 34 | 394228 | JAECYM000000000 | SAMN17025152 |
| c29VSb_15_M | vagina | healthy | 140 | 95 | O2:H5 | 5294433 | 82 | 220334 | JAECYL000000000 | SAMN17025153 |
| c31VSa_9_AN | vagina | healthy | 127 | - | O6:H31 | 5232112 | 66 | 388396 | JAECYK000000000 | SAMN17025154 |
| c32Ub_19_M | urine | rUTI | 12 | - | O4:H5 | 5095522 | 47 | 373088 | JAECYJ000000000 | SAMN17025155 |

A whole genome single nucleotide polymorphism (SNP) alignment of 11 sequenced strains is shown in Figure 2, together with antibiotic resistance phenotype. According to this phylogenetic tree, strains had ∼ 30000-49000 SNP differences, except strain belonging to ST73 (c13Ua_2_AN isolated from one rUTI donor) and its single locus variant (SLV) ST1154 (c11Ub_17_AN isolated from a healthy donor) with 1710 SNP differences. Two isolates belonging to ST131 (c14Ub_22_AE and c26Ub_7_AN) appear in a separate branch though they also differed in >13500 SNP. In fact, one of them (c14Ub_22_AE) exhibited *fimH41*-O25:H4 and the other one (c26Ub_7_AN) *fimH30-*O16:H5, suggesting they belong to different ST131 clades (see below). Antimicrobial susceptibility testing revealed that five isolates (Figure 2) were resistant to at least one antibiotic (amoxicillin-clavulanate acid, nalidixic acid, ciprofloxacin, tobramycin, gentamycin, tetracycline, trimethoprim or trimethoprim/sulfamethoxazole). One ST131 strain isolated from the urine of a healthy donor revealed multidrug resistance phenotype due to resistance to 6 antibiotics from 4 classes, including critical antibiotics for UTI treatment (trimethoprim/sulfamethoxazole and amoxicillin-clavulanate acid). ST12, ST140 (SLV 95) and ST127 strains were also resistant to amoxicillin-clavulanate, nalidixic acid and/or tetracycline (Figure 2). Detection of acquired resistance genes in our genomes was compatible with phenotypic characterization of resistance, with detection of *bla*_TEM-30_ in 4 strains resistant to amoxicillin-clavulanic acid combination, presence of *tet(A)* or *tet(B)* for tetracycline resistance strains or additional genes identified for multidrug resistant c26Ub_7_AN strain (e.g., *aac(3)-I id*, *aadA2*, *sul1*, *dfrA12*). We also observed mutations in intrinsic genes e.g., S83L mutation in *gyrA* gene for isolates c14Ub_22AE and c29VSb_15M, conferring resistance to fluoroquinolones, demonstrated previously by phenotypic antimicrobial susceptibility testing (AST) (Figure 2). We also detected a chromosomal mutation corresponding to resistance to fosfomycin, however their presence was not compatible with *in-vitro* AST results. Despite their phenotypic susceptibility to fosfomycin (Figure 2), in all 11 B2 strains we found E448K mutation in *glpT* gene, while in 2 ST131 strains we found additional E350Q in *uhpT* and V25I in *ptsI* genes. Although in *ptsI* gene besides well-known V25I mutation we found additional amino acid difference (R367K), the remaining hits were reported as strict prediction (above the bitscore cutoff).

**Figure 2.**
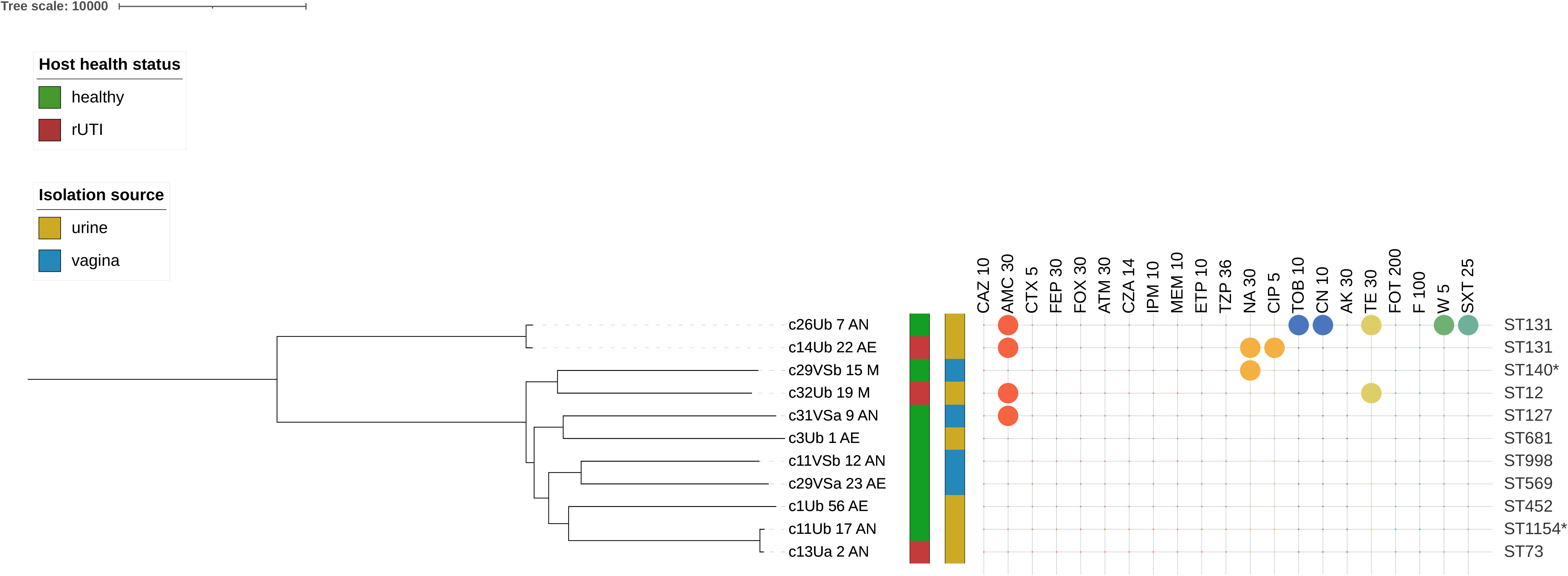
Phylogenetic tree representing whole genome SNPs alignment of 11 urogenital *E. coli* strains. Presence of colorful balls in the antibiotics chart represent resistance to certain antibiotics, while different colors represent different classes of antibiotics. An asterisk in the STs column means that this strain is single locus variant of another relative ST (see Table 2). *Antibiotics used: CAZ, ceftazidime; AMC, amoxicillin/clavulanic acid; CTX, cefotaxime; FEP, cefepime; FOX, cefoxitin; ATM, aztreonam; CZA, ceftazidime/avibactam; IPM, imipenem; MEM, meropenem; ETP, ertapenem; TZP, piperacillin/tazobactam; NA, nalidixic acid; CIP, ciprofloxacin; TOB, tobramycin; CN, gentamycin; AK, amikacin; TE, tetracycline; FOT, fosfomycin/trometamol; F, nitrofurantoin; W, trimethoprim; SXT, trimethoprim/sulfamethoxazole.

Screening of plasmid replicon sequences in these 11 isolates revealed that no known replicon sequences were detected for 4 strains (ST569, ST681, ST998, ST1154), whereas in the other B2 strains several replicon types were detected (**Supplementary Table S1**). Most of them are F-type plasmids (FII, FIA, FIB, FIC), being Col156 and IncFIB(AP001918) the most pervasive.

### Characterization and SNPs phylogeny of ST131 *E. coli*

In our cohorts (asymptomatic and rUTI women), we identified 2 isolates belonging to the pandemic *E. coli* clone ST131 that is responsible for a high rate of human UTI. To assess clade distribution and infer phylogeny of these isolates in relation to strain origin and host status, we performed SNP phylogenetic analysis with genomes available in National Center for Biotechnology Information (NCBI) public database for which metadata was available.

A total of 520 ST131 genomes was selected from NCBI public database, from which most were obtained from diseased human (D; n=422 strains) and much less from healthy carriers (H; n=98 strains). Regarding origin, the highest number of strains represented isolates of mixed origin, identified mainly in wound or pus, lung and sputum, blood or other bodily fluids (other; n=242 strains), followed by strains isolated from urogenital tract, specifically urine (n=179 strain) and gastrointestinal tract (n=99 strains). While isolates obtained from the gastrointestinal tract originated mostly from healthy human (H=96, D=3), isolates from urogenital were almost exclusively associated with urinary tract infections (H=1, D=178) and similarly, isolates from other human niches were mostly isolated from diseased host (H=1, D=241) e.g., bacteremia, sepsis, pneumonia.

The core genome phylogenetic tree (Figure 3), including 2 new genomes from this study and 3 reference strains, was congruent with previous ST131 phylogenetic inferences and establishes clustering of genomes in three clades corresponding to clade A (n=67), clade B (n=50) and clade C (n=406) (Figure 3). One of our ST131 strains (c14Ub_22_AE from rUTI woman) clustered in clade A (*fimH41*) and was highly similar to genomes of strains from urinary tract infection or gastrointestinal colonization showing only 32 or 30 SNP differences, respectively. Strain c26Ub_7_AN (from healthy woman), despite presenting *fimH30*, clustered in clade B isolates (carrying in most cases *fimH22* allele) and showed similarity to other clade B genomes from gastrointestinal tract colonizers (1111 SNPs) or infections (1157 SNPs). Interestingly, this is an unique representative of ST131 isolated from the urogenital tract deposited in NCBI that was obtained from a healthy individual.

**Figure 3.**
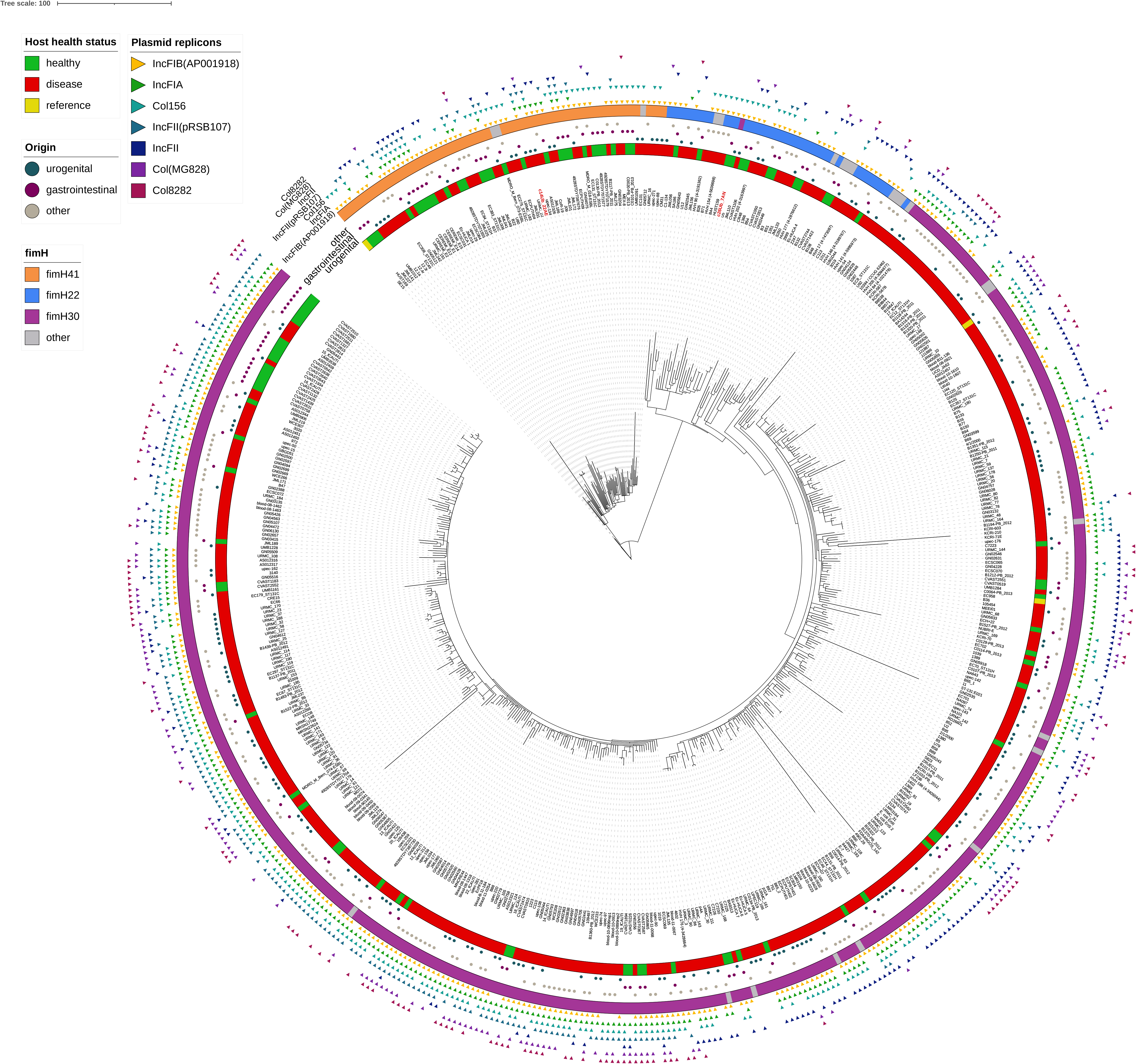
Core genome SNPs phylogenetic tree of 523 ST131 *E. coli* genomes of human origin. The alignment was performed using *E. coli* strain EC958 as a reference and *E. coli* SE15 was used as an outgroup. The metadata including host health status, strains origin, *fimH* type and presence of seven most common plasmid replicon sequences are incorporated in the tree, as explained in the legend. The identifiers of 2 strains from our urogenital collection are marked in red.

Overall, genomes isolated from healthy and diseased hosts were randomly distributed in the phylogeny, often with less than 50 SNPs among them. Similarly, isolates originated from urogenital tract or other body sites causing disease were highly similar to those colonizing either the gastrointestinal tract or the urogenital tract. SNPs matrix for ST131 core genome based phylogenetic tree is available in **Supplementary Table S2**.

Additionally, we detected 55 different plasmid replicon sequences however, only several were highly prevalent. Detection of seven most common (in minimum 25% of isolates) plasmid sequences i.e., IncFIB(AP001918), IncFIA, Col156, IncFII(pRSB107), IncFII, Col(MG828) and Col8282 is included in Figure 3. Among them, only IncFIA and IncFII were found to be significantly enriched in diseased isolates (p<0.05).

### Pangenome of ST131 *E. coli*

We performed pangenome analysis for *E. coli* ST131 collection, including 520 strains of human origin, with known isolation source and host health status. Pangenome was composed of 31089 genes, with only 2547 representing core genome (8.2%). Remaining genes represent accessory genome with 12295 being singletons (39.5%) i.e., specific to one genome only. Major part of pangenome are hypothetical proteins of unknown function (59.5%). Detailed list of all genes detected in pangenome analysis can be found in **Supplementary Table S3**.

We further investigated the accessory genome to clarify if the genomic content of the strains is anyhow specific to isolates obtained from different niches or from healthy or diseased host. We performed non-metric multidimensional scaling (NMDS) ordination based on 28542 genes matrix (accessory genome), applying additional metadata layers of origin and host health status (Figure 4). Although NMDS analysis demonstrated slight differences among some strains (tree different clusters observed represents A, B and C clades), there was obviously no distinction regarding isolates origin, and host health status, thus, the same conclusion can be drawn as from SNP analysis. Furthermore, we draw Venn Diagram to understand if there is any set of genes that could be specific to certain niche (Figure 5). In Figure 5 we can see that strains from each niche had plenty of shared genes, but they also had from 2384 to 7476 genes that have not been detected in isolates from other niches. We further investigated on the strains from the urogenital tract, what is the prevalence and type of those specific genes. Not surprisingly, 4215 genes (of 4711 total) represent singletons unique for only one genome. Only 496 genes were detected in more than one genome of urogenital origin, with the maximum prevalence of 8/500 genomes (1.5%). Moreover, most proteins detected do not have any predicted function (3383/4711, 71.8%). Among the known proteins, most are putatively involved in mobilome e.g., XerC tyrosin recombinase, various transposases or FlmA stable plasmid inheritance protein. Therefore, consistently with the previous conclusions, we did not identify any specific gene with significant prevalence that could be indicator of niche preferences. The list of all 4711 genes detected only among urogenital isolates is presented in **Supplementary Table S4**.

**Figure 4.**
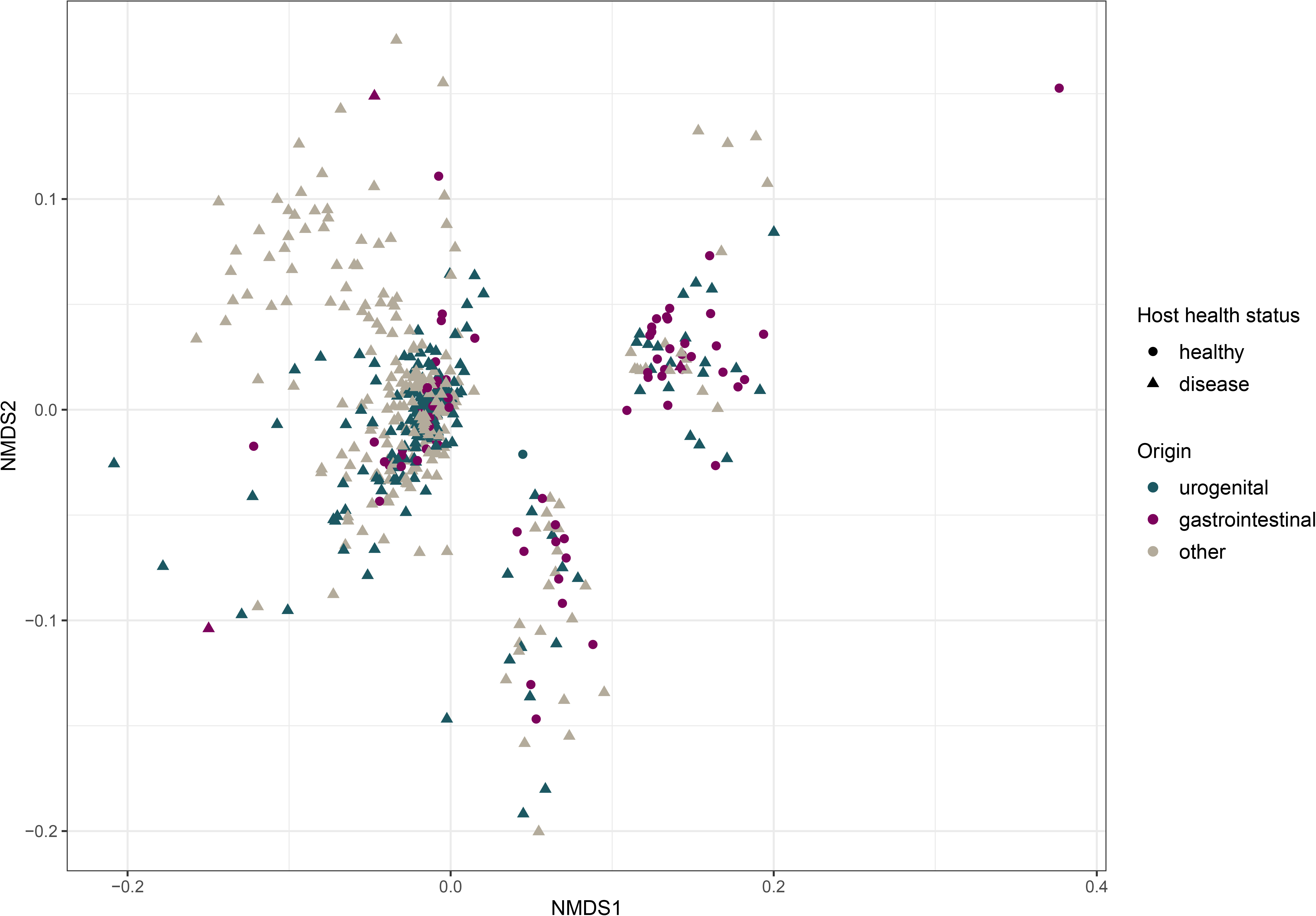
Non-metric multidimensional scaling (NMDS) based on 28542 genes matrix (accessory genome) extracted from 520 genomes. Metadata layers of origin and host health status are incorporated in the figure. Stress value for this ordination is 0.18.

**Figure 5.**
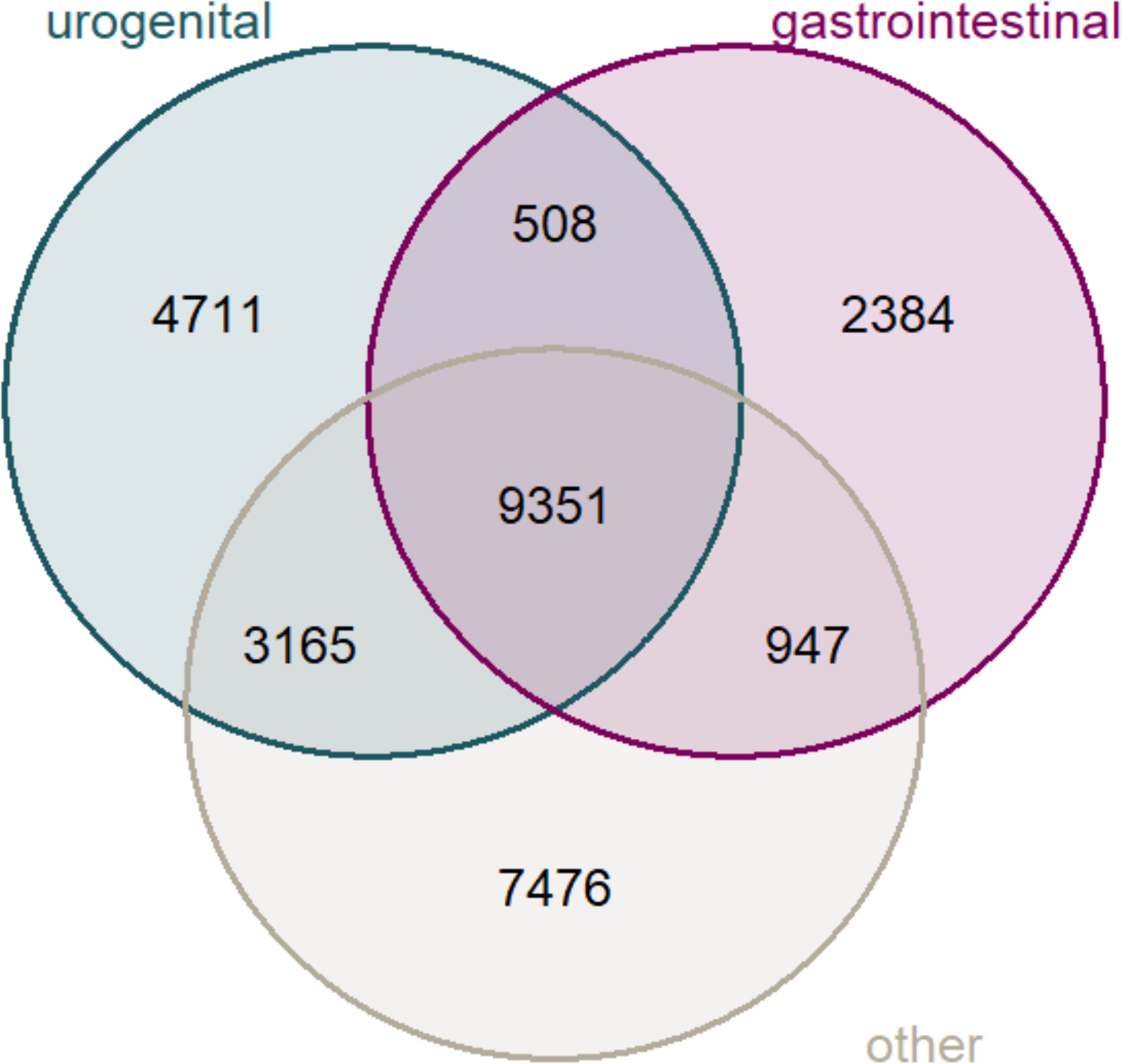
Venn Diagram for accessory genome (28542 genes) of 520 *E. coli* strains, representing sets according to strains isolation source i.e., urogenital (greenish), gastrointestinal (violet) and other (grey).

### Characterization and SNPs phylogeny of additional STs among urogenital *E. coli*

A whole genome comparison was performed between our B2 non-ST131 isolates and available genomes from the same ST on public databases, selected based also on host status and isolation source. In agreement with available literature, ST95 including ST140 (SLV 95) isolate (n=272) and ST73 including ST1154 (SLV 73) isolates (n=217) are the second and third most represented clonal groups (**Supplementary Figures S2 and S3,** respectively). SNP-based phylogenetic trees containing our ST140 (ST95 clonal complex), ST73 and ST1154 (both ST73 clonal complex) urogenital microbiome *E. coli* isolates also evidenced a high similarity between strains from different host status and origin, often showing <1000 SNPs differences. The same happened with the clones belonging to ST127 (n=66) and ST12 (n=45), that were closely related to genomes from isolates causing UTI and/or isolated from other niche (<300 SNP differences, **Supplementary Figures S4 and S5,** respectively).

The remaining 4 STs (ST452, ST569, ST681, ST998), all of them recovered from urinary microbiome of healthy women, were much less represented in NCBI database. Despite that, a comparative genomics analysis also showed for some of them a high similarity (from 390-9452 SNPs) to genomes deposited from other sources (Figure 6). Interestingly, although just few strains are available, they were isolated in different continents.

**Figure 6.**
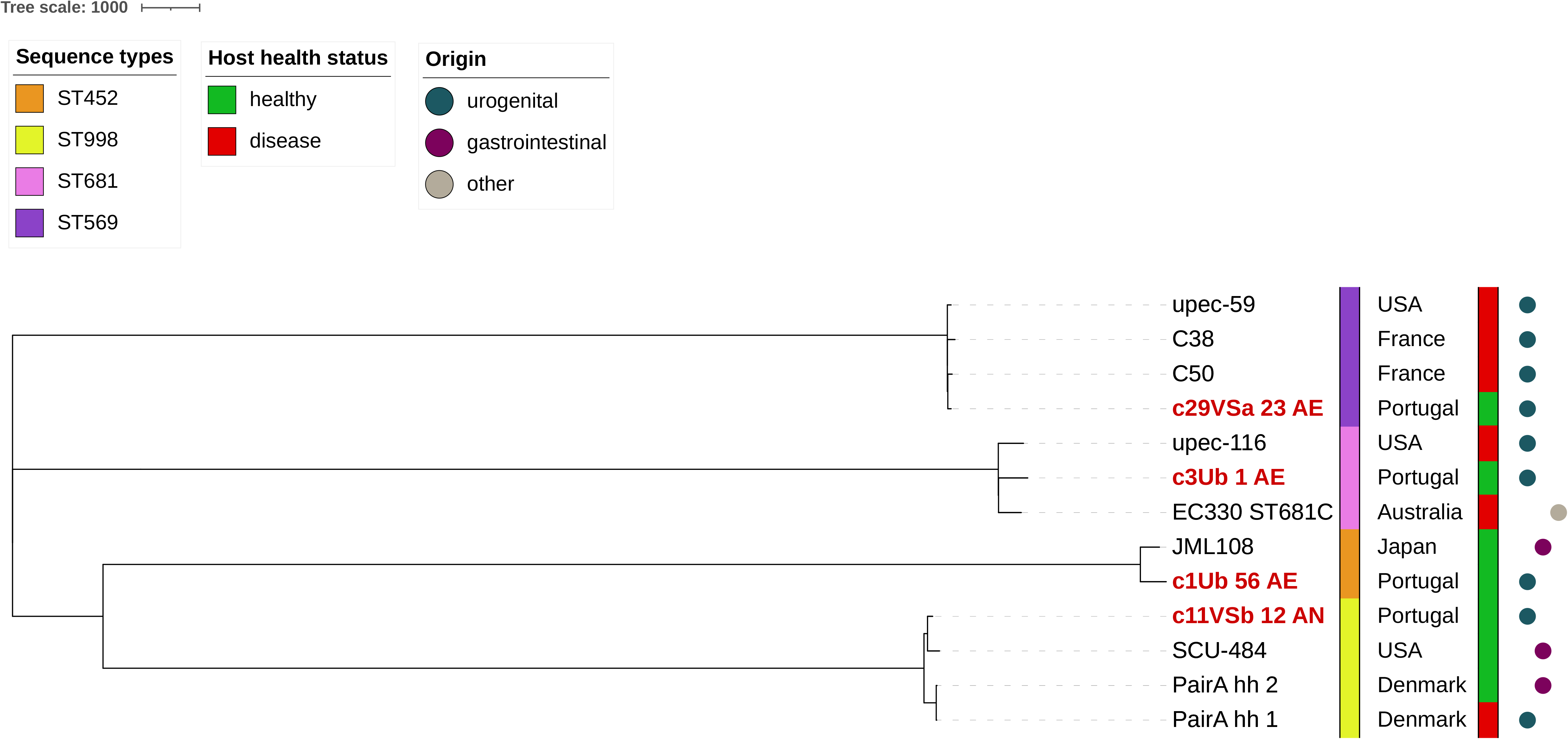
Whole genome SNPs phylogenetic tree of 4 infrequent STs (ST452, ST569, ST681, ST998) including 4 our urogenital strains (marked in red) and 9 genomes retrieved from NCBI database. The reference strain used for the alignment is *E. coli* 536. The tree is unrooted. Information on ST, health status of the host and origin of the strains is incorporated in the tree.

## DISCUSSION

In this study, we provided a snapshot on the *E. coli* strain diversity and genetic features of isolates identified in the female urogenital microbiome. We demonstrated a high occurrence of *E. coli* in the urogenital tract of asymptomatic women. Those *E. coli* were enriched in ExPEC strains particularly from the B2 phylogenetic group, frequently presenting resistance to at least one antibiotic commonly used to treat UTI and with similarity in genomic background and virulence gene profiles to those known to frequently cause UTI. Furthermore, we reported a ST131 strain with similar genetic background to those causing UTI inhabiting the urogenital tract of healthy woman.

*E. coli* was often found in both urine (29%) and vagina (37%) of healthy women, the latter revealing a prevalence slightly higher than that reported in previous studies^20, 33^. It is believed that there is an interconnectedness between these two niches^28^ in the same individual and in fact our data support that urine and vagina often share identical strains. However, in some samples, we also found variability in the number and type of *E. coli* strains identified in each of those niches, highlighting possible limitations in studies evaluating solely one of those niches for risk assessment of UTI acquisition. Furthermore, our data also highlights that strain level characterization is relevant to accurately evaluate contribution of these niches as reservoirs of strains with potential to persist and/or cause UTI.

We described a higher *E. coli* diversity in the urogenital microbiome of healthy women (phylogenetic groups A, B2, D, E, F) than that reported in the study from Garretto *et al.*, 2020 where only B2 *E. coli* strains (n=6) were identified from asymptomatic women^32^. In contrast to this ecological variation, only B2 *E. coli* strains were identified in our small set of samples from women with rUTI which is in line with the predominance of B2 strains in rUTI^34, 35^.

In the healthy cohort, we detected serogroups known to be commonly involved in UTI (e.g., O2, O6, O25)^13, 36, 37^, that together with previous report (3 urinary isolates of O2 and O6 serogroup^32^) points to a frequent carriage of those serogroups in healthy women urogenital microbiome. Remarkably, it is also of highlight the identification of ExPEC (n= 7 B2; n= 2 F) strains in > 50% of healthy women, enriched in putative virulence factors known to favor colonization and/or invasion of epithelial cells and development of UTI^6^. Similarly to previous studies, we showed that strains of B2 phylogenetic group possess wider range of putative virulence genes than other phylogenetic groups^16, 36, 38^. Moreover, virulence genes’ profiles detected in ExPEC isolates from healthy women were identical to those described in clinical isolates^16^. All other non-ExPEC strains except one phylogroup A *E. coli* possessed also several adhesins, siderophores and other putative virulence genes, likely supporting their adaptation and survival in the urogenital tract. Thus, in agreement with the study by Garretto *et al*., our study provides further evidence that urogenital tract carries strains with pathogenic potential.

Remarkably, many of our B2 strains (from either healthy or rUTI cohorts) belonged to worldwide disseminated *E. coli* lineages (ST12, ST73 complex, ST95 complex, ST131), that have been causing infections in humans^13, 15, 17, 18^. Our phylogenomics analysis revealed close relatedness with publicly available genomes regardless of human origin (host status or infection site), further demonstrating wide circulation of well-adapted and potentially pathogenic clones between different individuals.

The identification of strains, including ST131, in both cohorts that are resistant to antibiotic classes that are used to treat UTI (amoxicillin-clavulanate acid, ciprofloxacin, trimethoprim/sulfamethoxazole; Figure 2) represents an additional concern due to the risk of treatment failure. Moreover, antibiotic resistant strains in the microbiome constitute a reservoir of transferable antimicrobial resistance genes that can be shared with other strains/species by horizontal gene transfer^39, 40^. Thus, the possibility of enriching antibiotic resistant bacterial species (e.g., *E. coli*, *Citrobacter koserii*, *Klebsiella pneumoniae*)^29, 31^ in the urogenital microbiome can have a future direct implication into the human health.

Identification of ST131 in the urine of healthy woman challenges current understanding of the ecology of this pandemic clone. We demonstrate, for the first time, that the healthy urogenital microbiome can be a reservoir of putative pathogenic and antibiotic resistant ST131 strains. We demonstrated indistinguishable distribution of *E. coli* ST131 strains per origin (sample site and health status), providing further evidence that ST131 ExPEC causing infections in different human body sites and isolates from healthy urogenital tract are closely related, similarly to what was previously suggested for isolates from the gastrointestinal tract^41, 42^. This observation was further supported with NMDS analysis of 28542 accessory genes where no distinction was captured according to either isolates origin or host health status. Detection of *fimH30* allele in clade B isolates is not common but these recombination events were occasionally observed in other collections^11, 21^. Available data on prevalence of ST131 subclades is largely biased by human clinical isolates, and clade C in particular^43–45^. However, there is evidence that non-human sources are underestimated and there is a specialization for each clade, including a foodborne origin for clade B-*fimH22*^46^ and wastewater for clade A-*fimH41*^45^, suggesting those as additional possible sources of acquisition.

In this study, we also identified other intercontinental B2 *E. coli* clones i.e., ST95 complex, ST73 complex or ST127 from urogenital samples of healthy women. Furthermore, we performed MLST analysis on the B2 strains from asymptomatic women identified in the previous study by Garretto *et al*. and realized that those strains also belonged to intercontinental STs including ST95 (n=3), ST73 (n=1), ST12 (n=1) and ST1193 (n=1), all of them emerging global clones^21, 47, 48^. Altogether, these data suggest that healthy urogenital microbiome is a source of particularly widespread (e.g., food, domestic animals, environment) *E. coli* lineages that are frequently responsible for UTI^49, 50^.

Much less data is available on uncommon *E. coli* clones (ST452, ST569, ST681, ST998) detected in the urogenital microbiome of healthy women since they are poorly represented in public databases. Nevertheless, they had been previously reported as animal colonizers or from human infections in different continents (Figure 6), often associated with antibiotics resistance. For instance, *mcr-1* positive ST452 or ESBL producing ST998 *E. coli* strains were previously isolated from animals, including domestic animals i.e., dog or human UTI in five continents^51–55^. Similarly, ST681 strains were found nearly worldwide in animals (wild boars, non-human primates) and among isolates causing human ExPEC infections, including UTI^56–59^. Furthermore, besides reports on UTI caused by ST569, this lineage could have possible food and environmental reservoirs, where in the USA ST569 *E. coli* was found in meat^60^ or its detection in wastewater in South Africa^61^. Mbanga *et al*. also demonstrated that strain isolated from wastewater clustered together with previously isolated clinical ST998 *E. coli* isolated from UTI patient in the same area^52^. Interestingly, we found those strains only among healthy women and they were susceptible to all antibiotics tested.

In fact, the data and interpretation presented here, suggest that *E. coli* strains inhabiting the urogenital tract of healthy women are frequently ExPEC intercontinental lineages, likely due to their widespread and, consequently, a higher probability of human exposure. Nevertheless, whether those strains possess the ability to persistently colonize human urogenital tract or they are only transient microbiome members remains to be evaluated. Furthermore, one of the most fundamental questions is still unanswered, specifically, what are the circumstances that would trigger infection development by those colonizing strains.

We recognize as limitations of this study the small sample size and the absence of fecal samples from the same individuals to assess concomitant gastrointestinal colonization, however, *in silico* comparative genomic analysis performed show close relatedness of urinary and gastrointestinal strains. Using voided urine has been also considered a limitation due to the possibility of collecting microbes that reside in the vaginal environment, but we demonstrated that voided urine sample can provide distinct strains than those found in vagina. Furthermore, insights from this type of sample are valuable as it is used in current UTI diagnostic practices.

## CONCLUSIONS

In this study we demonstrated that *E. coli* strains identified in the urogenital microbiome of healthy women belong frequently to international, pandemic, and occasionally antibiotic resistant B2 clones, further supporting the role of the urogenital tract as a source of ExPEC strains prone to cause UTI. These findings are a hallmark for further understanding the ecology of these clones and their distribution in human host reservoirs.

The influence of microbial communities, host and environmental features triggering the transition from colonization to infection is still unknown, but this subject deserves further investigation.

## MATERIAL AND METHODS

### Study design and sample description

*Escherichia coli* strains were isolated in the framework of a study on Female Urogenital Microbiome conducted at Faculty of Pharmacy, University of Porto, Portugal (2016-2019)^31^. All women provided informed written consent for participation in the study. The study was developed according to the Helsinki Declaration principles and the protocol was submitted and approved by the Ethical Commission of Faculty of Pharmacy, University of Porto. This study included isolates obtained from: i) 21 healthy women (aged 24-57) that were asymptomatic for any urinary disorder and that declared to be in good health condition, from which urogenital samples (31 urine and 30 vaginal swabs) were analyzed (10 women provided samples twice^31^); and ii) isolates from 3 women with rUTI from which only urinary isolates were studied (3 urine samples). Two of rUTI women were in remission phase (no current manifestation of UTI) when the sample was collected (donor 13 and 14; **Supplementary Table S1**), and remaining (donor 32; **Supplementary Table S1**) provided a sample during UTI. Of note, latter donor was likely colonized with the same *E. coli* clone during remission prior to this subsequent UTI (strains similarity assessed by PFGE and 41 virulence associated genes profile; data not shown).

### Identification of *E. coli* isolates

All samples were analyzed based on extended culturomic protocol previously published^31^. From a total of 64 samples analyzed (34 voided urine and 30 vaginal swabs), mean of 3 isolates/sample (up to 8 colonies/sample; proportionally to the bacterial load) were selected with morphology characteristic for *E. coli*.

Isolates were preliminarily identified by MALDI-TOF MS (VITEK MS, bioMérieux, France), and further confirmed by amplification of *malB* gene^62^. A total of 65 *E. coli* isolates was identified and stored, of which 22 isolates originated from urine of healthy donors (1-7/sample), 28 from vaginal swabs of healthy donors (1-5/sample) and 15 from urine of donors with history of rUTI (1-8/sample).

### Characterization of *E. coli* population

Identification of *E. coli* phylogenetic groups and virulence gene profiling were performed on all *E. coli* isolates to discard overrepresented genotypes per sample. *E. coli* phylogenetic groups were identified according to the revisited method proposed by Clermont et al.^63^. The presence of 41 VFs associated with ExPEC pathotype including adhesins, toxins, siderophores, capsule type, protectins, invasins and miscellaneous was screened by PCR (**Supplementary Table S5**). Detection of *hlyA* operon duplication was performed by PCR, as previously described^64^ (**Supplementary Table S5**). The strains were classified as putative ExPEC if fulfilled following criteria: presence of >2 amongst *papAH* and/or *papC*, *sfa/focDE*, *afa/draBC*, *kpsM II* and *iutA*^65^. Based on phylogenetic groups screening and VF profiles including 41 genes we selected 25 representative urinary and vaginal *E. coli* strains that are described in detail in **Supplementary Table S1**.

When *E. coli* isolates from the same individual but different sample type (urine and vaginal strains) presented identical VF profile, PFGE was used to further assess relationships of isolates originating from the same donor. The standard protocol for PFGE adapted from Gautom et al.^66^ was used with 1.6% chromosomal grade agarose (SeaKem® Gold Agarose) for plugins preparation, 1.2% gel with pulse-field certified agarose (SeaKem® Gold Agarose) for PFGE run, genomic DNA digested with 20U *XbaI* and CHEF DNA Size Standard – Lambda Ladder used as size marker. The running conditions included initial switch time of 2.2 seconds, final switch time of 63.8 seconds and run time of 19 hours with 6 V/cm and included angle of 120.

Based on PFGE fingerprints, for the purpose of further characterization of urogenital *E. coli* population, we discarded overrepresented isolates from the same women (the same strain in urine and vagina), and we identified 19 strains representing unique genotypic profiles, independently of sample type.

*In vitro* antibiotic susceptibility testing was evaluated by disk diffusion for 21 antibiotics (diverse β-lactams, fluoroquinolones, aminoglycosides, tetracyclines, fosfomycin, nitrofurantoin, trimethoprim, sulfamethoxazole-trimethoprim), according to EUCAST guidelines (www.eucast.org).

### Whole genome sequencing

Considering that B2 *E. coli* strains represented 11/19 (58%) of representative isolates from both healthy and rUTI cohorts and revealed ExPEC pathotype features, these strains were characterized by whole genome sequencing (WGS) and comparative genomics approaches, as described below. In case of identical isolates from the same donor estimated previously by PFGE, the vaginal isolate was discarded, and urinary isolate was subjected to WGS.

Genomic DNA of 11 representative strains from healthy (n=4 urine, n=4 vaginal) and rUTI (n=3 urine) cohorts belonging to B2 phylogenetic group was extracted (Wizard Genomic DNA Purification Kit, PROMEGA) and sequenced by Illumina NovaSeq 2×150 nt. Reads were trimmed by Trimmomatic^67^ version 0.39 and quality of reads checked by FastQC version 0.11.9 (http://www.bioinformatics.babraham.ac.uk/projects/fastqc/). *De novo* assembly was performed by SPAdes^68^ version 3.13.0 and quality of assembly checked by Quast^69^ version 5.0.2. Annotations of the draft genomes were provided by the NCBI Prokaryotic Genome Annotation Pipeline, and additionally by Prokka^70^ version 1.14.6.

The Whole Genome Shotgun project has been deposited at DDBJ/ENA/GenBank under the BioProject accession number PRJNA548360. Accession numbers to WGS submission and BioSample per each strain are available in Table 2.

Characterization of strains including serotype, *E. coli* phylogenetic group and multilocus sequence typing (Achtman scheme) was assessed using available *in silico* tools (http://www.genomicepidemiology.org/; http://clermontyping.iame-research.center/). Virulence profile of each strain previously defined by PCR method was verified and extended to 49 putative virulence genes using in-house databases and ncbi-blast-2.8.1+ package (https://ftp.ncbi.nlm.nih.gov/blast/executables/blast+/LATEST/). Antimicrobial resistance genes were identified using ResFinder 4.1 (https://cge.cbs.dtu.dk/services/ResFinder/) and Resistance Gene Identifier within The Comprehensive Antibiotic Resistance Database (https://card.mcmaster.ca/analyze/rgi), plasmid replicon sequences by PlasmidFinder 2.1 (https://cge.cbs.dtu.dk/services/PlasmidFinder/) and *fimH* typing using FimTyper 1.0 (https://cge.cbs.dtu.dk/services/FimTyper/).

### Comparative genomics of B2 *E. coli* clones

Genomes from B2 strains sequenced in this study were compared with genomes deposited on public databases according to the clone identified by MLST. The NCBI Assembly public database was assessed at 14/07/2020 and 19668 *E. coli* assemblies were downloaded, including 1266 complete genomes and 18402 draft genomes (7,432 scaffolds and 10,970 contigs). Additional 66 assemblies of *E. coli* of urinary origin published in 09/2020 by Garretto et al.^32^, potentially relevant for this study and not available previously, were retrieved from NCBI Assembly database. In total, 19734 assemblies were subjected to mlst 2.19.0 pipeline^71^ (https://github.com/tseemann/mlst) to access STs of available strains. Strains representing STs detected among our strains i.e., ST131, ST95, ST73, ST12, ST140, ST452, ST569, ST681, ST998 and ST1154 were extracted. Only genomes with available metadata were considered to allow selection of those isolated from humans and with available information on host health status and isolation source. Following these criteria, we grouped isolates according to host health status (healthy-H or diseased-D), and origin (urogenital tract, gastrointestinal tract or other isolation source). Additionally, genomes well known as references for certain ST were included in the analysis.

For the purpose of this manuscript, based on available epidemiological data, we divided described STs in two following groups: intercontinental STs including ST131, ST95 clonal complex, ST73 clonal complex, ST12 and ST127; and uncommon STs including ST452, ST569, ST681 and ST998.

### Single nucleotide polymorphisms analysis

Selected STs were subjected to SNP-based phylogenetic analysis. The genomes of specific STs were subjected to snippy (version 4.4.0) variant calling pipeline (https://github.com/tseemann/snippy) using snippy-multi and mapping to appropriate reference genome (stated in description of the figures). A core genome and whole genome SNPs alignments generated by snippy were used for Gubbins pipeline (Genealogies Unbiased By recomBinations In Nucleotide Sequences)^72^ version 2.4.1 to remove recombinant regions. The SNP matrices from core genome and whole genome alignment for comparison was generated with snp-dist version 0.7.0 (https://github.com/tseemann/snp-dists). The resulting SNP-based alignments were used to reconstruct phylogeny for each ST using RAxML, with appropriate outgroups. The phylogenetic trees obtained were represented with Interactive Tree of Life (iTOL, https://itol.embl.de).

### Pangenome analysis

For pangenome analysis of ST131 *E. coli* strains 520 genomes were included representing strains only of human origin, with known host health status and isolation source. Genomes were annotated with prokka version 1.14.6, and pangenome analysis was performed with Roary^73^ version 3.13.0 with the default criteria for proteins identity of 95%. Genes were represented by following collections: core genes were present in 99-100% genomes, accessory genes were present in less than 99% of genomes, including singletons, that are unique for one genome only. The data was reshaped and clean with tidyverse^74^ package version 1.3.0. Further analysis including NMDS and calculation of stress value was performed in R^75^ version 4.0.3 with package vegan^76^ version 2.5.7 and ggplot2^77^ version 3.3.5. Venn Diagram was performed using VennDiagram package^78^ version 1.6.20. Proteins specific to each origin group were also identified with VennDiagram package.

### Statistical analysis

Continuous variables were interpreted based on descriptive statistics. Welch Two Sample t-test in R^75^ version 3.6.2 was used to access significance of phylogenetic groups and VF distribution between the different sample types and cohorts. Heatmap and clustering of the isolates based on VFs presence/absence was performed with Pheatmap package version 1.0.12 (https://github.com/raivokolde/pheatmap), with clustering based on Euclidean distance.

## AUTHOR STATEMENTS

### Authors’ contributions

M.K. performed all experimental and computational work (genomic extraction and analysis, comparative genomics, pangenome analysis, antibiotic susceptibility testing) and wrote the manuscript. A.N. supervised the study design, contributed to data interpretation, revised and edited the manuscript. L.P. contributed to reviewing the manuscript, project design and administration, and funding.

## Acknowledgements

We would like to thank Svetlana Ugarcina Perovic for performing MALDI-TOF MS analysis and bioMérieux (Portugal, Lda) for MALDI-TOF MS equipment and material provided. We also would like to acknowledge Svetlana Ugarcina Perovic, Joana Rocha and Filipa Grosso fundamental contribution to Female Urogenital Microbiome Project.

## ETHICS DECLARATIONS

### Ethical statement

Approval of the study was obtained from the Faculty of Pharmacy (University of Porto, Porto, Portugal) Ethics Committee. Procedures performed in the study were all in accordance with the ethical standards of the institutional and national research committee, with the 1964 Helsinki Declaration, and its later amendments. All individual participants included in the study had given written informed consent.

## DATA AVAILABLILITY STATEMENT

The Whole Genome Shotgun project of the isolates c1Ub_56_AE, c3Ub_1_AE, c11Ub_17_AN, c11VSb_12_AN, c13Ua_2_AN, c14Ub_22_AE, c26Ub_7_AN, c29VSa_23_AE, c29VSb_15_M, c31VSa_9_AN, c32Ub_19_M have been deposited at DDBJ/ENA/GenBank under the accession numbers JAECYT000000000, JAECYS000000000, JAECYR000000000, JAECYQ000000000, JAECYP000000000, JAECYO000000000, JAECYN000000000, JAECYM000000000, JAECYL000000000, JAECYK000000000, JAECYJ000000000, respectively.

## ABBREVIATIONS

AST: antimicrobial susceptibility testing
CFU: colony forming units
EUCAST: The European Committee on Antimicrobial Susceptibility Testing
ExPEC: extraintestinal pathogenic *Escherichia coli*
FUM: female urogenital microbiome
MALDI-TOF MS: Matrix-assisted laser desorption/ionization time-of-flight mass spectrometry
NCBI: National Center for Biotechnology Information
NMDS: Non-metric multidimensional scaling
PCR: polymerase chain reaction; PFGE, Pulsed-field Gel Electrophoresis
rUTI: recurrent urinary tract infection
SLV: Single locus variant, SNP, single nucleotide polymorphism
ST: Sequence type
U: urine
UTI: urinary tract infection
WGS: whole genome sequencing
VF: virulence factor
VS: vaginal swab.

## REFERENCES

1. Negus, M., Phillips, C. & Hindley, R. Recurrent urinary tract infections: a critical review of the currently available treatment options. Obstet Gynaecol 22, 115–121 (2020).

2. Öztürk, R. & Murt, A. Epidemiology of urological infections: a global burden. World J Urol 38, 2669–2679 (2020).

3. Subashchandrabose, S. & Mobley, H. L. T. Virulence and Fitness Determinants of Uropathogenic *Escherichia coli*. Microbiol Spectr 3, (2015).

4. Foxman, B. The epidemiology of urinary tract infection. Nat Rev Urol 7, 653–660 (2010).

5. Medina, M. & Castillo-Pino, E. An introduction to the epidemiology and burden of urinary tract infections. Ther Adv Urol 11, (2019).

6. Flores-Mireles, A. L., Walker, J. N., Caparon, M. & Hultgren, S. J. Urinary tract infections: epidemiology, mechanisms of infection and treatment options. Nat Rev Microbiol 13, 269– 284 (2015).

7. Tamadonfar, K. O., Omattage, N. S., Spaulding, C. N. & Hultgren, S. J. Reaching the End of the Line: Urinary Tract Infections. Microbiol Spectr 7, (2019).

8. Russo, T. A. & Johnson, J. R. Proposal for a new inclusive designation for extraintestinal pathogenic isolates of *Escherichia coli*: ExPEC. J Infect Dis 181, 1753–1754 (2000).

9. Manges, A. R. et al. Global Extraintestinal Pathogenic *Escherichia coli* (ExPEC) Lineages. Clin. Microbiol. Rev. 32, (2019).

10. Brzuszkiewicz, E. et al. How to become a uropathogen: comparative genomic analysis of extraintestinal pathogenic *Escherichia coli* strains. Proc Natl Acad Sci U S A 103, 12879– 12884 (2006).

11. Matsumura, Y. et al. Rapid Identification of Different *Escherichia coli* Sequence Type 131 Clades. Antimicrob Agents Chemother 61, (2017).

12. Stoesser, N. et al. Evolutionary History of the Global Emergence of the *Escherichia coli* Epidemic Clone ST131. mBio 7, e02162 (2016).

13. Ciesielczuk, H. et al. Trends in ExPEC serogroups in the UK and their significance. Eur. J. Clin. Microbiol. Infect. Dis. 35, 1661–1666 (2016).

14. Kallonen, T. et al. Systematic longitudinal survey of invasive *Escherichia coli* in England demonstrates a stable population structure only transiently disturbed by the emergence of ST131. Genome Res 27, 1437–1449 (2017)

15. Fibke, C. D. et al. Genomic Epidemiology of Major Extraintestinal Pathogenic *Escherichia coli* Lineages Causing Urinary Tract Infections in Young Women Across Canada. Open Forum Infect Dis 6, ofz431 (2019).

16. Flament-Simon, S.-C. et al. Clonal Structure, Virulence Factor-encoding Genes and Antibiotic Resistance of *Escherichia coli*, Causing Urinary Tract Infections and Other Extraintestinal Infections in Humans in Spain and France during 2016. Antibiotics (Basel) 9, (2020).

17. Johnson, J. R. et al. Household Clustering of *Escherichia coli* Sequence Type 131 Clinical and Fecal Isolates According to Whole Genome Sequence Analysis. Open Forum Infect Dis 3, ofw129 (2016).

18. Banerjee, R. & Johnson, J. R. A new clone sweeps clean: the enigmatic emergence of *Escherichia coli* sequence type 131. Antimicrob Agents Chemother 58, 4997–5004 (2014).

19. Lewis, A. L. & Gilbert, N. M. Roles of the vagina and the vaginal microbiota in urinary tract infection: evidence from clinical correlations and experimental models. GMS Infect Dis 8, (2020).

20. O’Brien, V. P. et al. Low-dose inoculation of *Escherichia coli* achieves robust vaginal colonization and results in ascending infection accompanied by severe uterine inflammation in mice. PLoS One 14, e0219941 (2019).

21. Johnson, J. R. & Russo, T. A. Molecular Epidemiology of Extraintestinal Pathogenic *Escherichia coli*. EcoSal Plus 8, (2018).

22. Brannon, J. R. et al. Invasion of vaginal epithelial cells by uropathogenic *Escherichia coli*. Nat Commun 11, 2803 (2020).

23. Neugent, M. L., Hulyalkar, N. V., Nguyen, V. H., Zimmern, P. E. & Nisco, N. J. D. Advances in Understanding the Human Urinary Microbiome and Its Potential Role in Urinary Tract Infection. mBio 11, (2020).

24. Jones-Freeman, B. et al. The microbiome and host mucosal interactions in urinary tract diseases. Mucosal Immunol 14 779–792 (2021).

25. Siddiqui, H., Nederbragt, A. J., Lagesen, K., Jeansson, S. L. & Jakobsen, K. S. Assessing diversity of the female urine microbiota by high throughput sequencing of 16S rDNA amplicons. BMC Microbiol. 11, 244 (2011).

26. Wolfe, A. J. et al. Evidence of uncultivated bacteria in the adult female bladder. J. Clin. Microbiol. 50, 1376–1383 (2012).

27. Price, T. K. et al. The Clinical Urine Culture: Enhanced Techniques Improve Detection of Clinically Relevant Microorganisms. J Clin Microbiol 54, 1216–1222 (2016).

28. Thomas-White, K. et al. Culturing of female bladder bacteria reveals an interconnected urogenital microbiota. Nat Commun 9, (2018).

29. Coorevits, L., Heytens, S., Boelens, J. & Claeys, G. The resident microflora of voided midstream urine of healthy controls: standard versus expanded urine culture protocols. Eur. J. Clin. Microbiol. Infect. Dis. 36, 635–639 (2017).

30. Khasriya, R. et al. Spectrum of bacterial colonization associated with urothelial cells from patients with chronic lower urinary tract symptoms. J. Clin. Microbiol. 51, 2054–2062 (2013).

31. Ksiezarek, M., Ugarcina-Perovic, S., Rocha, J., Grosso, F. & Peixe, L. Long-term stability of the urogenital microbiota of asymptomatic European women. BMC Microbiol 21, 64 (2021).

32. Garretto, A. et al. Genomic Survey of *E. coli* From the Bladders of Women With and Without Lower Urinary Tract Symptoms. Front Microbiol 11, (2020).

33. Salinas, A. M. et al. Vaginal microbiota evaluation and prevalence of key pathogens in ecuadorian women: an epidemiologic analysis. Sci Rep 10, 18358 (2020).

34. Luo, Y. et al. Similarity and divergence of phylogenies, antimicrobial susceptibilities, and virulence factor profiles of *Escherichia coli* isolates causing recurrent urinary tract infections that persist or result from reinfection. J Clin Microbiol 50, 4002–4007 (2012).

35. Karami, N., Lindblom, A., Yazdanshenas, S., Lindén, V. & Åhrén, C. Recurrence of urinary-tract infections with extended-spectrum β-lactamase-producing *Escherichia coli* caused by homologous strains among which clone ST131-O25b is dominant. J Glob Antimicrob Resist 22, 126–132 (2020).

36. Johnson, J. R. & Stell, A. L. Extended virulence genotypes of *Escherichia coli* strains from patients with urosepsis in relation to phylogeny and host compromise. J. Infect. Dis. 181, 261–272 (2000).

37. Sarkar, S., Ulett, G. C., Totsika, M., Phan, M.-D. & Schembri, M. A. Role of capsule and O antigen in the virulence of uropathogenic *Escherichia coli*. PLoS One 9, e94786 (2014).

38. Vejborg, R. M., Hancock, V., Schembri, M. A. & Klemm, P. Comparative Genomics of *Escherichia coli* Strains Causing Urinary Tract Infections. Appl Environ Microbiol 77, 3268–3278 (2011).

39. Liu, L. et al. The human microbiome: a hot spot of microbial horizontal gene transfer. Genomics 100, 265–270 (2012).

40. Langille, M. G. I., Meehan, C. J. & Beiko, R. G. Human microbiome: a genetic bazaar for microbes? Curr Biol 22, R20–22 (2012).

41. Nielsen, K. L., Dynesen, P., Larsen, P. & Frimodt-Møller, N. Faecal *Escherichia coli* from patients with *E. coli* urinary tract infection and healthy controls who have never had a urinary tract infection. J Med Microbiol 63, 582–589 (2014).

42. Moreno, E. et al. Relationship between *Escherichia coli* strains causing urinary tract infection in women and the dominant faecal flora of the same hosts. Epidemiol Infect 134, 1015–1023 (2006).

43. . Petty, N. K., et al. Global dissemination of a multidrug resistant *Escherichia coli* clone. Proc Natl Acad Sci U S A 111, 5694–5699 (2014).

44. Peirano, G. et al. Characteristics of *Escherichia coli* Sequence Type 131 Isolates That β-Lactamases: Global Distribution of the H30-Rx Sublineage. Antimicrob. Agents Chemother 58, 3762–3767 (2014).

45. Finn, T. J. et al. A Comprehensive Account of *Escherichia coli* Sequence Type 131 in Wastewater Reveals an Abundance of Fluoroquinolone-Resistant Clade A Strains. Appl Environ Microbiol 86, e01913–19 (2020).

46. Liu, C. M. et al. *Escherichia coli* ST131-H22 as a Foodborne Uropathogen. mBio 9, (2018).

47. Valenza, G. et al. First report of the new emerging global clone ST1193 among clinical β-lactamase (ESBL)-producing *Escherichia coli* from Germany. J Glob Antimicrob Resist 17, 305–308 (2019).

48. Ding, Y., Zhang, J., Yao, K., Gao, W. & Wang, Y. Molecular characteristics of the new emerging global clone ST1193 among clinical isolates of *Escherichia coli* from neonatal invasive infections in China. Eur J Clin Microbiol Infect Dis 40, 833–840 (2021).

49. Riley, L. W. Extraintestinal Foodborne Pathogens. Annu Rev Food Sci Technol 11, 275–294 (2020).

50. Manges, A. R. & Johnson, J. R. Reservoirs of Extraintestinal Pathogenic *Escherichia coli*. Microbiol Spectr 3, (2015).

51. Delgado-Blas, J. F., Ovejero, C. M., Abadia-Patiño, L. & Gonzalez-Zorn, B. Coexistence of mcr-1 and blaNDM-1 in *Escherichia coli* from Venezuela. Antimicrob. Agents Chemother 60, 6356–6358 (2016).

52. Mbelle, N. M. et al. The Resistome, Mobilome, Virulome and Phylogenomics of Multidrug-Resistant *Escherichia coli* Clinical Isolates from Pretoria, South Africa. Sci Rep 9, 16457 (2019).

53. Bozcal, E., Eldem, V., Aydemir, S. & Skurnik, M. The relationship between phylogenetic classification, virulence and antibiotic resistance of extraintestinal pathogenic *Escherichia coli* in İzmir province, Turkey. PeerJ 6, e5470 (2018).

54. Alghoribi, M. F. et al. Antibiotic-resistant ST38, ST131 and ST405 strains are the leading uropathogenic *Escherichia coli* clones in Riyadh, Saudi Arabia. J Antimicrob Chemother 70, 2757–2762 (2015).

55. Yasugi, M. et al. Whole-genome analyses of extended-spectrum or AmpC β-lactamase producing Escherichia coli isolates from companion dogs in Japan. PLoS One 16, e0246482 (2021).

56. Alonso, C. A., González-Barrio, D., Ruiz-Fons, F., Ruiz-Ripa, L. & Torres, C. High frequency of B2 phylogroup among non-clonally related fecal *Escherichia coli* isolates from wild boars, including the lineage ST131. FEMS Microbiol Ecol 93, (2017).

57. Foster-Nyarko, E. et al. Genomic diversity of *Escherichia coli* isolates from non-human primates in the Gambia. Microb Genom 6, (2020).

58. Hertz, F. B. et al. Population structure of Drug-Susceptible, -Resistant and ESBL-producing *Escherichia coli* from Community-Acquired Urinary Tract Infections. BMC Microbiol 16, 63 (2016).

59. Banerjee, R. et al. The Clonal Distribution and Diversity of Extraintestinal *Escherichia coli* Isolates Vary According to Patient Characteristics. Antimicrob. Agents Chemother. 57, 5912–5917 (2013).

60. Yamaji, R. et al. A Population-Based Surveillance Study of Shared Genotypes of Escherichia coli Isolates from Retail Meat and Suspected Cases of Urinary Tract Infections. mSphere 3, (2018).

61. Mbanga, J. et al. Genomic Insights of Multidrug-Resistant *Escherichia coli* From Wastewater Sources and Their Association With Clinical Pathogens in South Africa. Front Vet Sci 8, 636715 (2021).

62. Wang, R. F., Cao, W. W. & Cerniglia, C. E. PCR detection and quantitation of predominant anaerobic bacteria in human and animal fecal samples. Appl Environ Microbiol 62, 1242– 1247 (1996).

63. Clermont, O., Christenson, J. K., Denamur, E. & Gordon, D. M. The Clermont *Escherichia coli* phylo-typing method revisited: improvement of specificity and detection of new phylo-groups. Environ Microbiol Rep 5, 58–65 (2013).

64. Ksiezarek, M. et al. Phylogenomic analysis of a highly virulent *Escherichia coli* ST83 lineage with potential animal-human transmission. Microb Pathog 155, 104920 (2021).

65. Johnson, J. R. et al. Isolation and Molecular Characterization of Nalidixic Acid-Resistant Extraintestinal Pathogenic *Escherichia coli* from Retail Chicken Products. Antimicrob Agents Chemother 47, 2161–2168 (2003).

66. Gautom, R. K. Rapid pulsed-field gel electrophoresis protocol for typing of *Escherichia coli* O157:H7 and other gram-negative organisms in 1 day. J Clin Microbiol 35, 2977–2980 (1997).

67. Bolger, A. M., Lohse, M. & Usadel, B. Trimmomatic: a flexible trimmer for Illumina sequence data. Bioinformatics 30, 2114–2120 (2014).

68. Bankevich, A. et al. SPAdes: a new genome assembly algorithm and its applications to single-cell sequencing. J. Comput. Biol. 19, 455–477 (2012).

69. Gurevich, A., Saveliev, V., Vyahhi, N. & Tesler, G. QUAST: quality assessment tool for genome assemblies. Bioinformatics 29, 1072–1075 (2013).

70. Seemann, T. Prokka: rapid prokaryotic genome annotation. Bioinformatics 30, 2068–2069 (2014).

71. Jolley, K. A. & Maiden, M. C. BIGSdb: Scalable analysis of bacterial genome variation at the population level. BMC Bioinformatics 11, 595 (2010).

72. Croucher, N. J. et al. Rapid phylogenetic analysis of large samples of recombinant bacterial whole genome sequences using Gubbins. Nucleic Acids Res 43, e15 (2015).

73. Page, A. J. et al. Roary: rapid large-scale prokaryote pan genome analysis. Bioinformatics 31, 3691–3693 (2015).

74. Wickham, H. et al. Welcome to the Tidyverse. Journal of Open Source Software 4, 1686 (2019).

75. R Core Team. R: A language and environment for statistical computing. R Foundation for Statistical Computing, Vienna, Austria. https://www.R-project.org/ (2018).

76. Oksanen, J. et al. vegan: Community Ecology Package. https://CRAN.R-project.org/package=vegan (2019).

77. Wickham, H. ggplot2: Elegant Graphics for Data Analysis. Springer-Verlag New York (2016).

78. Chen, H. & Boutros, P. C. VennDiagram: a package for the generation of highly-customizable Venn and Euler diagrams in R. BMC Bioinformatics 12, 35 (2011).

