## Supplementary Figures for "The darkest place is under the candlestick - healthy urogenital tract as a source of UTI-related *Escherichia coli* lineages"

**Supplementary Figure S1**. Pulsed-field Gel Electrophoresis (PFGE) of selected E. coli isolates.

**M** - marker; **1** - 1Ub_83; **2** - 1VSb_14; **3** - 10Ua_105; **4** - 10VSa_39; **5** - 15Ub_26; **6** - 15VSb_20; **7** - 29VSa_23; **8** - 29VSb_15; **9** - 3Ub_1; **10** - 3VSb_22; **11** - 11Ub_17; **12** - 11VSb_7; **13** - 11VSb_12; **14** - 26Ub_7; **15** - 26VSb_8; **-** other isolates.

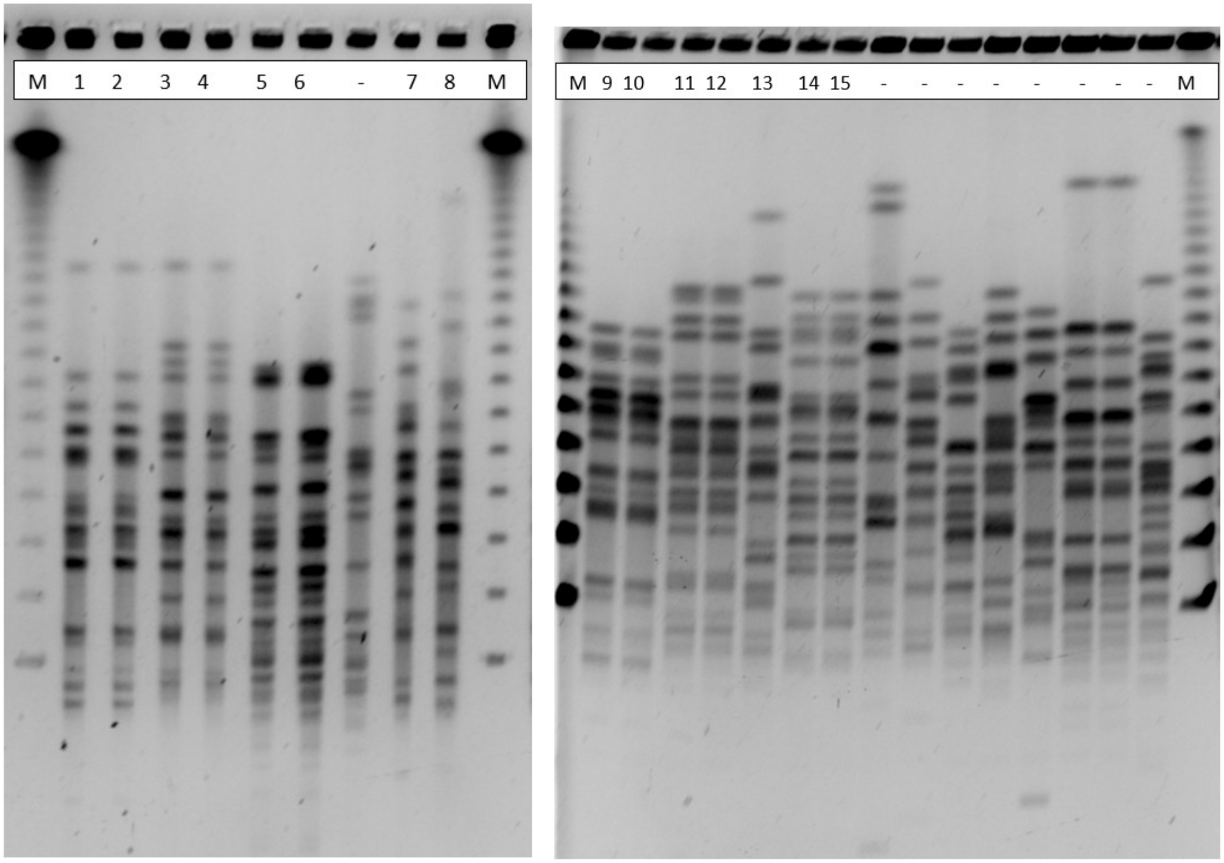

**Supplementary Figure S2.** Whole genome SNPs phylogenetic tree of 272 ST95 and ST140 (SLV 95) *E. coli* genomes of human origin. The alignment was performed using *E. coli* strain UTI89 as a reference. The metadata including host health status and strains origin is incorporated in the tree, as explained in the legend. The identifier of 1 strain from our urogenital collection is marked in red. The identifiers of 3 strains from Garretto et al. 2020 are marked in blue.

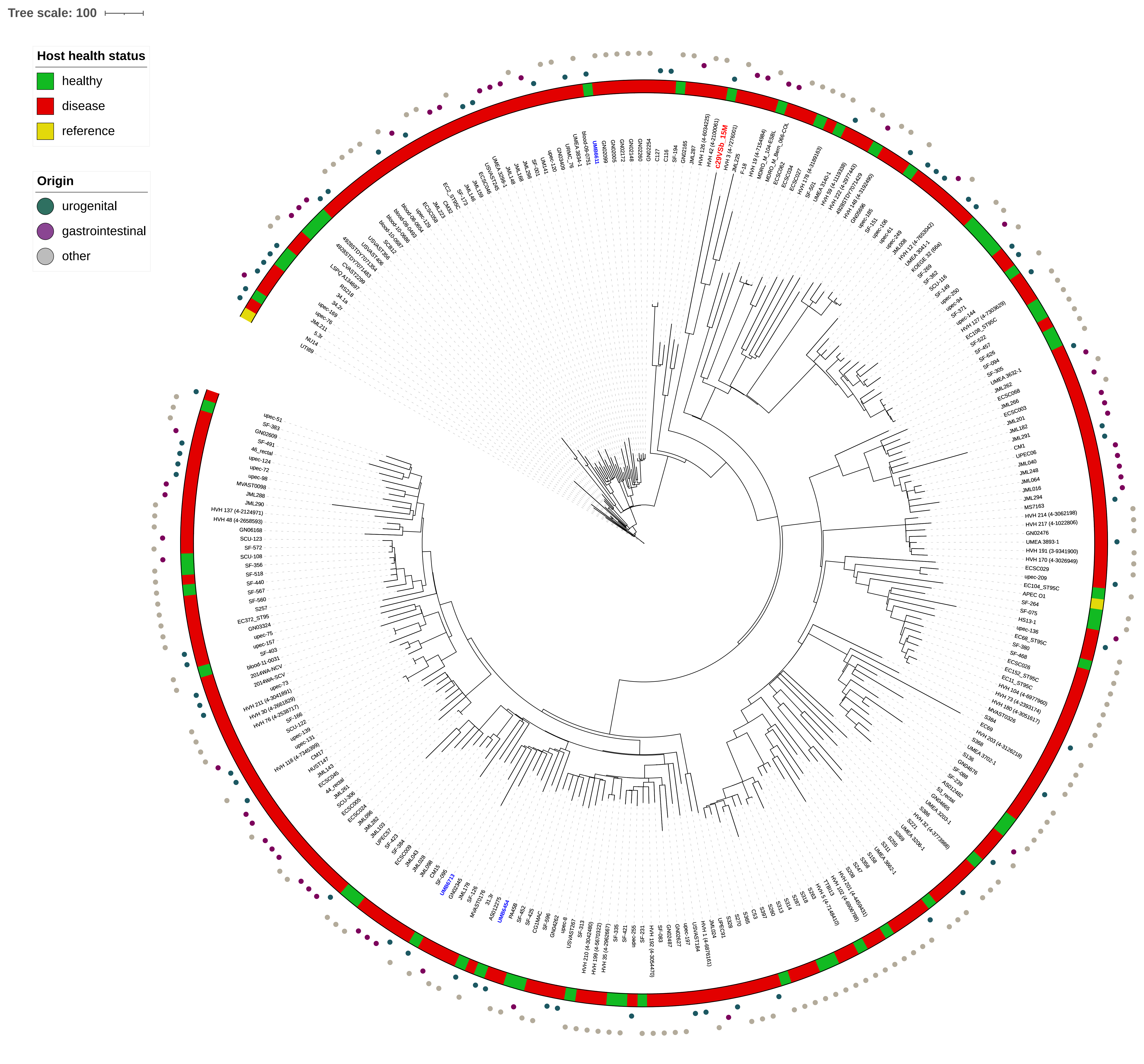

Tree scale: 100
Host health status
healthy
disease
reference
Origin
urogenital
gastrointestinal
other
c29VSb_15M

**Supplementary Figure S3.** Whole genome SNPs phylogenetic tree of 217 ST73 and ST1154 (SLV 73) *E. coli* genomes of human origin. The alignment was performed using *E. coli* strain CFT073 as a reference. The metadata including host health status and strains origin is incorporated in the tree, as explained in the legend. The identifiers of 2 strain from our urogenital collection are marked in red.

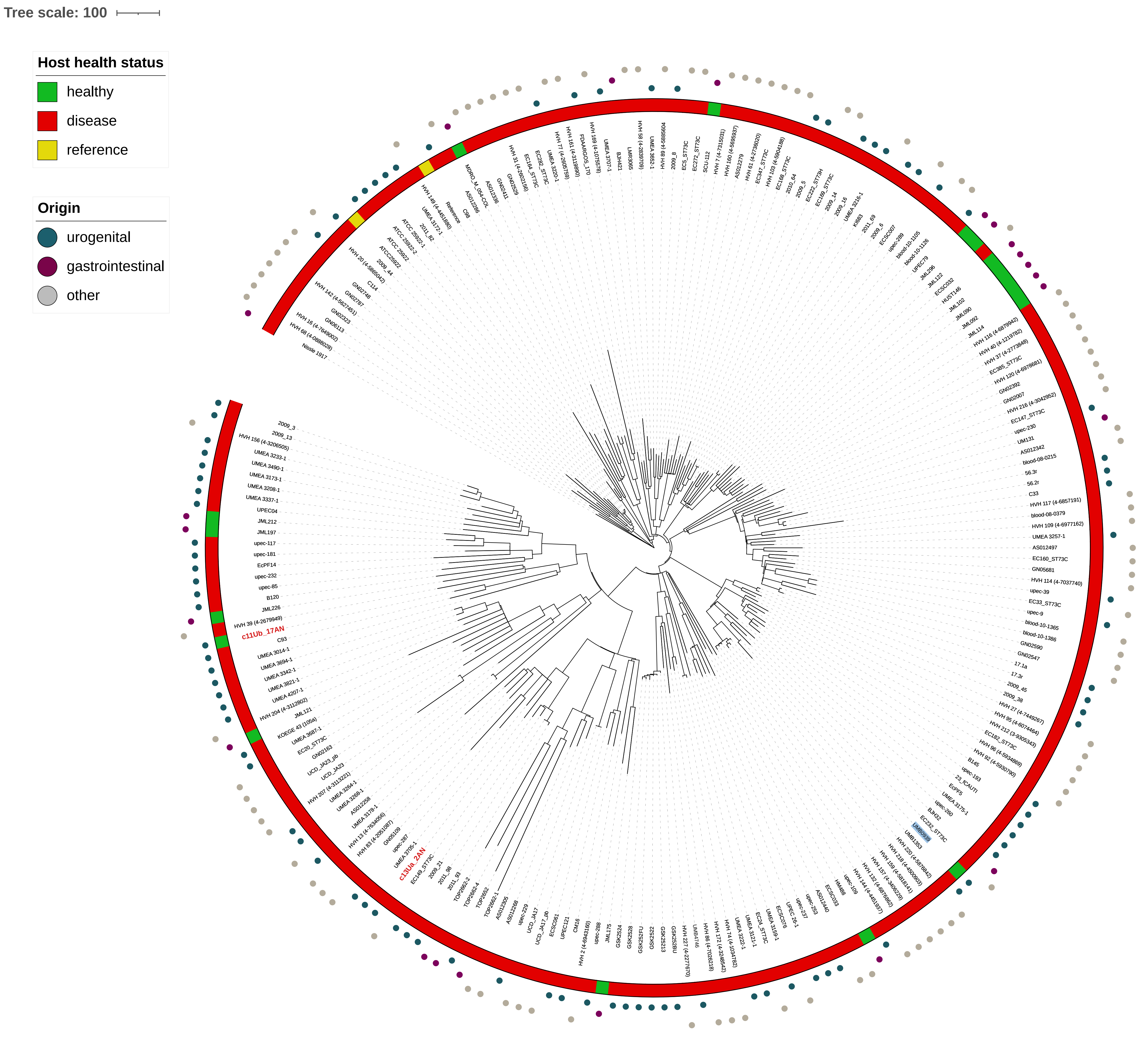
Tree scale: 100
Host health status
healthy
disease
reference
Origin
urogenital
gastrointestinal
other
c11Ub_17AN
c13Ua_2AN

**Supplementary Figure S4.** Whole genome SNPs phylogenetic tree of 66 ST127 *E. coli* genomes of human origin. The alignment was performed using *E. coli* strain 536 as a reference. The metadata including host health status and strains origin is incorporated in the tree, as explained in the legend. The identifier of strain from our urogenital collection is highlighted in green.

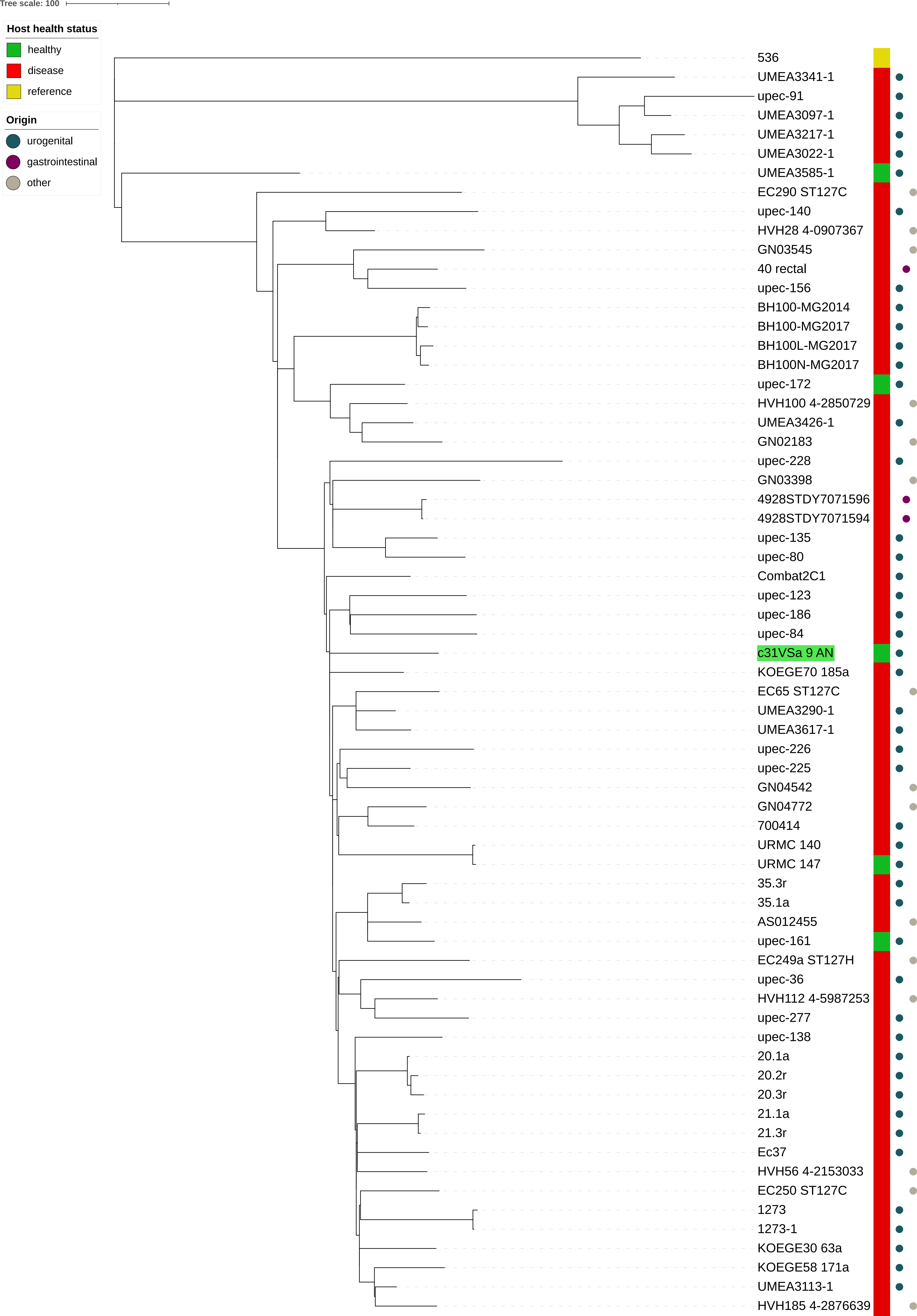
Tree scale: 100
Host health status
healthy
disease
reference
Origin
urogenital
gastrointestinal
other
536
UMEA3341-1
upec-91
UMEA3097-1
UMEA3217-1
UMEA3022-1
UMEA3585-1
EC290 ST127C
upec-140
HVH28 4-0907367
GN03545
40 rectal
upec-156
BH100-MG2014
BH100-MG2017
BH100L-MG2017
BH100N-MG2017
upec-172
HVH100 4-2850729
UMEA3426-1
GN02183
upec-228
GN03398
4928STDY7071596
4928STDY7071594
upec-135
upec-80
Combat2C1
upec-123
upec-186
upec-84
c31VSa 9 AN
KOEGE70 185a
EC65 ST127C
UMEA3290-1
UMEA3617-1
upec-226
upec-225
GN04542
GN04772
700414
URMC 140
URMC 147
35.3r
35.1a
AS012455
upec-161
EC249a ST127H
upec-36
HVH112 4-5987253
upec-277
upec-138
20.1a
20.2r
20.3r
21.1a
21.3r
Ec37
HVH56 4-2153033
EC250 ST127C
1273
1273-1
KOEGE30 63a
KOEGE58 171a
UMEA3113-1
HVH185 4-2876639

**Supplementary Figure S5.** Whole genome SNPs phylogenetic tree of 45 ST12 *E. coli* genomes of human origin. The alignment was performed using *E. coli* strain D8 as a reference. The metadata including host health status and strains origin is incorporated in the tree, as explained in the legend. The identifier of strain from our urogenital collection is highlighted in green.

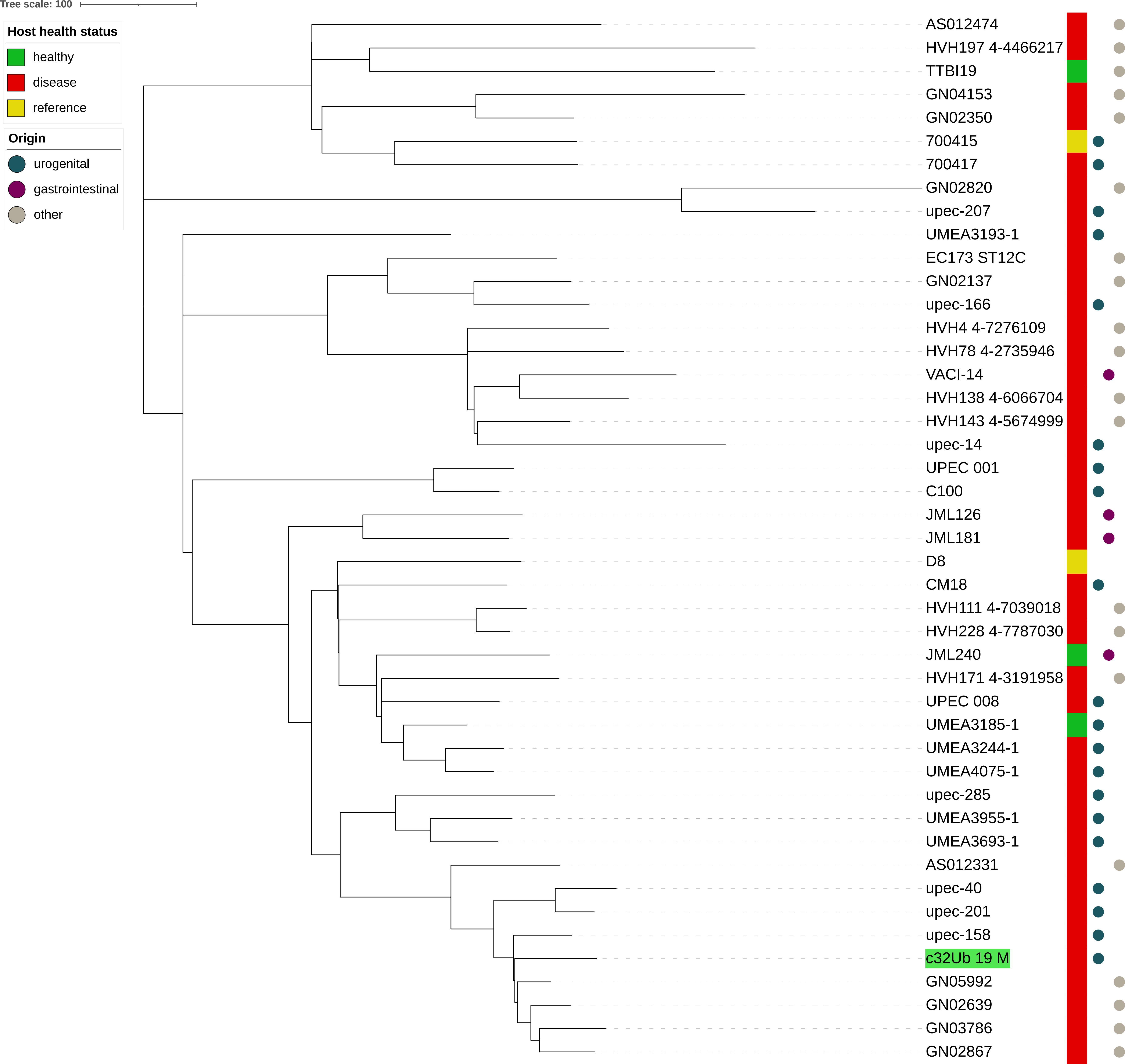
Tree scale: 100
Host health status
healthy
disease
reference
Origin
urogenital
gastrointestinal
other
AS012474
HVH197 4-4466217
TTBI19
GN04153
GN02350
700415
700417
GN02820
upec-207
UMEA3193-1
EC173 ST12C
GN02137
upec-166
HVH4 4-7276109
HVH78 4-2735946
VACI-14
HVH138 4-6066704
HVH143 4-5674999
upec-14
UPEC 001
C100
JML126
JML181
D8
CM18
HVH111 4-7039018
HVH228 4-7787030
JML240
HVH171 4-3191958
UPEC 008
UMEA3185-1
UMEA3244-1
UMEA4075-1
upec-285
UMEA3955-1
UMEA3693-1
AS012331
upec-40
upec-201
upec-158
c32Ub 19 M
GN05992
GN02639
GN03786
GN02867
